# Circadian Rhythms are Disrupted in Patients and Preclinical Models of Machado-Joseph Disease

**DOI:** 10.1101/2025.01.03.631212

**Authors:** Rodrigo F.N. Ribeiro, Dina Pereira, Sara M. Lopes, Tiago Reis, Patrick Silva, Diana D. Lobo, Laetitia S. Gaspar, João Durães, Ana Rita Fernandes, Marisa Ferreira-Marques, Catarina Carvalhas-Almeida, João Peça, Ana Rita Álvaro, Isabel Santana, Magda M. Santana, Maria Manuel C. Silva, Luís Pereira de Almeida, Cláudia Cavadas

**Affiliations:** Center for Neuroscience and Cell Biology (CNC-UC), Univ. of Coimbra, Coimbra, Portugal, 3004-504; Center for Innovation in Biomedicine and Biotechnology (CIBB), Univ. of Coimbra, Coimbra, Portugal; Faculty of Pharmacy, Univ. of Coimbra (FFUC), Coimbra, Portugal, 3000-548; Gene Therapy Center of Excellence (GeneT), Coimbra, Portugal, 3004-504; Department of Life Sciences, Faculty of Sciences and Technology, University of Coimbra, 3000-456, Coimbra, Portugal; Institute for Interdisciplinary Research, Univ. of Coimbra (iiiUC), Coimbra, Portugal, 3030-789; Department of Neurology, Coimbra Hospital and University Centre (CHUC), ULS de Coimbra, Portugal, 3004-561; Faculty of Medicine, Univ. of Coimbra (FMUC), Coimbra, Portugal, 3000-548

**Keywords:** Machado-Joseph Disease, Circadian rest-activity rhythms, Spinocerebellar ataxia type 3, Core body temperature, Suprachiasmatic nucleus, Core clock genes

## Abstract

Machado-Joseph disease (MJD) is caused by an abnormal CAG repeat expansion in the *ATXN3* gene, leading to the expression of a mutant ataxin-3 (mutATXN3) protein. MJD patients exhibit a wide range of clinical symptoms, including motor incoordination. Emerging evidence highlights circadian rhythm disruptions as early indicators and potential risk factors for the progression of neurodegenerative conditions. Circadian rhythms are regulated by internal clocks, with the suprachiasmatic nucleus (SCN) acting as the master pacemaker to synchronize timing across the body’s behavioural and physiological functions. While sleep disturbances have been observed in MJD, the role of clock regulation in its pathophysiology remains largely unexplored in spinocerebellar ataxias. This study aimed to investigate circadian rhythms, characterize associated disruptions, and uncover the mechanisms underlying clock dysregulation in patients and preclinical models of MJD.

Circadian activity in MJD patients was assessed over two weeks using actigraphy, while in a YAC-MJD transgenic mouse model, circadian rhythms were examined through: (a) wheel-running experiments; (b) telemetry-based monitoring of core body temperature; (c) immunohistochemical analysis of the neuropeptides arginine vasopressin (AVP) and vasoactive intestinal polypeptide (VIP) in the SCN and paraventricular nucleus (PVN); and (d) RT-qPCR evaluation of clock gene expression in the cerebellum. The impact of mutATXN3 on clock mechanisms was further investigated using *Bmal1*/*Per2-*luciferase reporters.

MJD patients exhibited a progressive decline in robustness of behavioural rhythms, demonstrated by negative correlations between the circadian function index, rest-activity fragmentation, and sleep efficiency with MJD clinical scales. YAC-MJD mice exhibited reduced activity levels, increased behavioural fragmentation, and required three additional days to re-entrain after a jet lag protocol, compared to controls. Disrupted core body temperature rhythms were observed, including a phase advance and elevated temperature (∼1 °C) at the onset of the active period. Furthermore, transgenic mice showed reduced levels of VIP and AVP in the SCN and PVN, and decreased clock gene expression in the cerebellum. Lastly, we found new mechanistic evidence that WT ATXN3 activates the promoters of *Bmal1* and *Per2*, whereas mutATXN3 loses the capacity to drive *Per2* upon polyglutamine expansion.

Overall, our findings indicate that central clock dysfunction in MJD is associated with impaired clock gene expression and disruptions in activity and temperature rhythms. This study provides the first robust evidence of circadian rhythm dysregulation and underlying mechanisms in MJD, paving the way for the identification of new biomarkers and the development of novel circadian-based interventions to tackle MJD and possibly other spinocerebellar ataxias.

**Graphical abstract:** 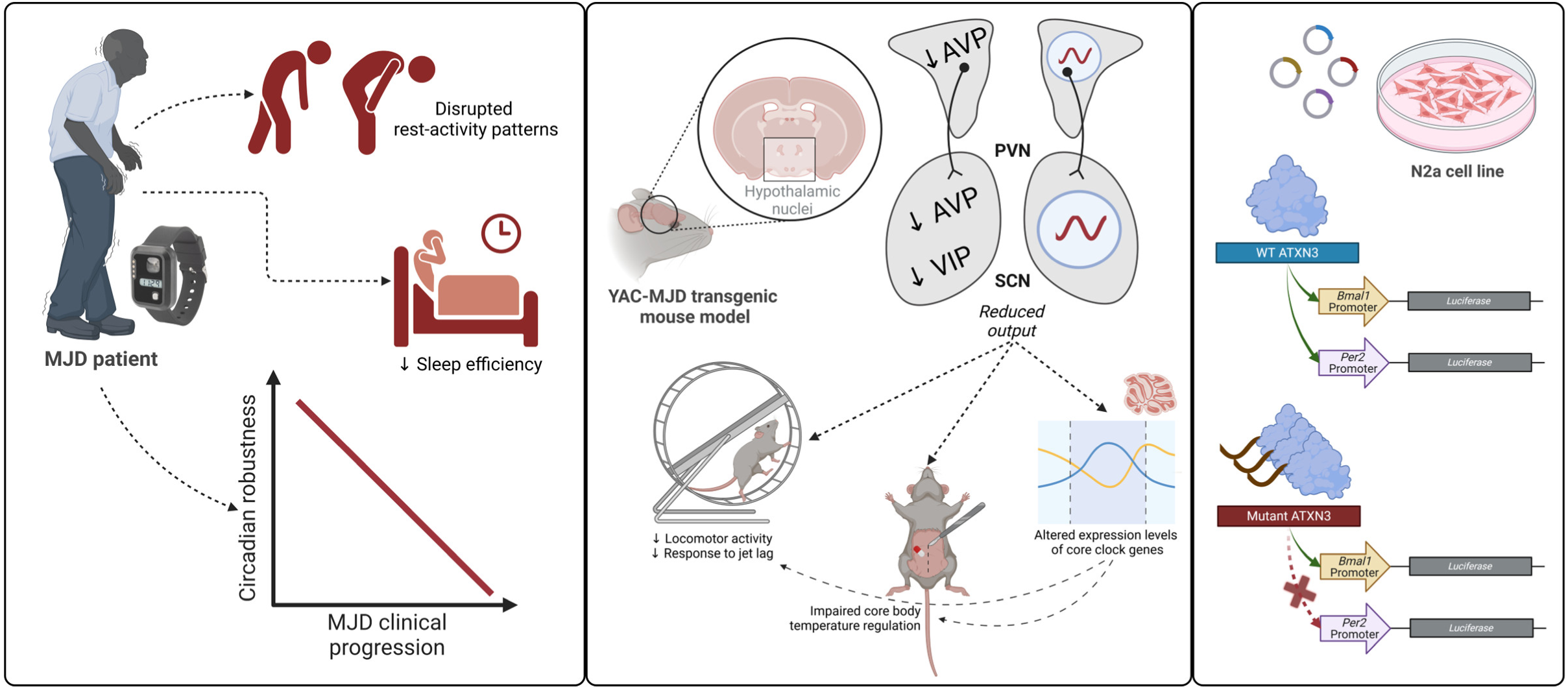

## Introduction

Machado-Joseph disease (MJD), also known as spinocerebellar ataxia type 3 (SCA3), is the most frequent autosomal dominantly inherited ataxia worldwide.^1^ MJD is caused by a mutation in the *ATXN3* gene, located on chromosome 14q32.12, that consists of an abnormal CAG repeat expansion ^2–4^. This mutation results in the expression of an expanded polyglutamine tract in the ataxin-3 protein (ATXN3) that becomes prone to aggregation, leading to neurodegeneration in several brain regions, mainly in the cerebellum.^3,5,6^ MJD manifests at an adult age with a wide range of clinical symptoms, including motor incoordination, gait and limb ataxia, dystonia, distal muscular atrophy, dysarthria, and ocular abnormalities.^7–9^ Currently, only symptomatic treatments are available for patients with MJD, none of which is able to cure or reduce the progression of this incapacitating and fatal disease.^10,11^ Furthermore, while sleep problems have been described in MJD ^12–14^, the role of circadian rhythms in its pathophysiology remains largely unexplored, both in this condition and in other spinocerebellar ataxias (SCAs).

Circadian rhythms are driven by a hierarchically organized timing system composed of a central master clock (the suprachiasmatic nucleus – SCN), located in the hypothalamus, and peripheral clocks, located in virtually all cells. Together, these biological clocks regulate multiple molecular, cellular, physiological and behavioural processes, including body temperature, hormone release, heart rate, sleep-wake cycles, and locomotion.^15–19^ Circadian rhythms depend on cell-autonomous and autoregulatory transcriptional/translational feedback loops of clock genes that oscillate on a 24 h cycle.^15,16,19^ These networks involve key components of the positive limb — *Brain and muscle ARNT-like protein* (*BMAL1/2*), *Circadian locomotor output cycles kaput* (*CLOCK*), and *RAR-related orphan receptor* (*RORα/β/c*) — which activate the transcription of negative limb players, including *Period* (*PER1/2/3*) and *Cryptochrome* (*CRY1/2*), that feedback to inhibit the activity of the positive limb.^15,16,18^ In the mammalian circadian timekeeping system, the SCN is at the top of the hierarchy and is required for the synchronization of the peripheral clocks throughout the body and the coordination of their functions.^15,16,18–20^ For that, a robust intercellular synchronization within the SCN, driven by arginine vasopressin (AVP) and vasoactive intestinal polypeptide (VIP), is crucial for sustaining the strength of downstream oscillators via neuronal and neuroendocrine pathways.^18,20^ Importantly, the disruption of the central clock and circadian rhythms has been linked to several pathologies.^16,17,19^

Despite differences in the pathogenesis of neurodegenerative disorders such as Alzheimer’s disease, Parkinson’s disease, and Huntington’s disease, a common hallmark among them is the disruption of circadian rhythms, which is thought to contribute to both disease onset and clinical profile.^17^ These dysfunctions include disturbances in sleep-wake cycles, rest-activity rhythms, core body temperature (CBT), clock gene expression, melatonin and cortisol levels, SCN firing and reduced levels of its synchronizers, AVP and VIP, as previously reviewed.^17,21–24^ Notably, the physiological consequences of circadian and sleep dysfunctions rank among the most debilitating symptoms experienced by patients with neurodegenerative conditions.^17,25^

While circadian disruptions are well-documented in other neurodegenerative diseases, their role in SCAs, including MJD, remains less explored. Emerging evidence suggests that the circadian system may play a significant role in the pathophysiology of MJD. Firstly, restless leg syndrome and obstructive sleep apnea have been found to be prevalent in MJD patients.^12–14^ Secondly, a dysregulated sleep architecture and altered EEG spectral power band were observed in a mouse model of the disease.^26^ Additionally, *Atxn3* expression follows an oscillatory profile over a 24 h period ^27^, and *Bmal1* knock-out mice exhibit reduced levels of *Atxn3* in the cerebellum.^28^ Another important observation is the decreased expression of RORα in a mouse model of MJD^29^, with RORα knock-down mice showing cerebellar ataxia.^30^ We previously found that MJD patients exhibit reduced levels of SIRT1, a NAD^+^-dependent deacetylase essential for clock regulation^17^, and that restoring this protein in a mouse model of the disease significantly improved neuropathology and motor coordination.^11^ Notably, other studies have shown that mice lacking SIRT1 in the brain exhibit several circadian rhythm alterations, while SIRT1 overexpression protects against circadian dysfunction.^31^

In this context, this study aimed to investigate circadian rhythms, characterize associated disruptions and uncover the mechanisms underlying clock dysregulation in patients and preclinical models of MJD. For this purpose, we conducted a clinical study to assess the rest-activity rhythms of MJD patients and found several progressive circadian alterations. To understand the neurobiological mechanisms behind this dysfunction, we analysed a YAC-MJD transgenic mouse model, widely used in MJD preclinical studies as it mirrors many of its pathological characteristics.^32^ We found reduced and more fragmented activity levels, impaired light cycle re-entrainment, and a disrupted CBT rhythm. Our findings suggest that mutant ATXN3 (mutATXN3) may induce alterations in the regulation of clock genes and key neurons within the central clock, such as those in the SCN and PVN, thereby contributing to the observed circadian dysfunctions. Furthermore, we show that both WT ATXN3 and mutATXN3 can drive the promoter of *Bmal1*, but only the wild-type form effectively promotes the transcription of *Per*2, highlighting a novel mechanism regulating the clock and how circadian rhythms may be disrupted in MJD pathology.

These findings open avenues for the development of biomarkers and therapeutic strategies targeting circadian rhythms to improve patient outcomes in this devastating disease.

## Materials and methods

### Study participants and data collection

This study included genetically confirmed MJD patients who were followed up at the Department of Neurology, Coimbra Hospital and University Centre (CHUC) and the Center for Neuroscience and Cell Biology, University of Coimbra (CNC-UC). The study protocol was approved by the local ethics committee and all participants provided informed consent prior to enrolment, in accordance with the Declaration of Helsinki. Demographic information (sex and age) and clinical data (scale scores, disease duration, and *ATXN3* genotype determined by PCR-based fragment length analysis) were collected. To ensure consistency and minimize potential biases, data collection followed a standardized protocol that included a comprehensive interview, a clinical and functional evaluation, and questionnaires. Clinical evaluation included the Scale for the Assessment and Rating of Ataxia (SARA) and the Inventory of Non-Ataxic Signs (INAS). SARA scale was used to assess performance in items associated with cerebellar ataxia, the most relevant and predominant alteration in MJD.^33,34^ INAS was used to determine the presence of non-ataxia signs.^35^ Functional performance was assessed using the Composite Cerebellar Functional Severity Score (CCFS) test and the Activities of Daily Living (ADL) scale. CCFS was used to address cerebellar ataxia in the upper limbs^36^, while ADL scale allowed the assessment of the quality of life of the patients and their ability to live independently.^37^ Sleep quality was assessed using the Pittsburgh Sleep Quality Index (PSQI) questionnaire.^38^ Patients were classified according to the SARA score as mildly ataxic (3.0 ≤ SARA ≤ 7.5), moderately ataxic (8.0 ≤ SARA ≤ 23.5), or severely ataxic (SARA ≥ 24.0).

For the monitoring of rest-activity cycles, all patients were asked to wear the ActTrust 2 actigraphy device (Condor Instruments, SP, Brazil) on the non-dominant wrist for 2 weeks under normal ambulatory conditions. During this period, patients were instructed to maintain their usual lifestyle and record their sleep patterns in a sleep diary. The device was programmed to continuously record motor activity (using the Proportional Integral Mode, PIM), skin temperature (°C), and light exposure (lux) at one-minute intervals (epoch). PIM is known to have a stronger correlation with polysomnography. It assesses movement intensity based on the area under the signal curve for each epoch.^39^ Raw actigraphy data were processed using the algorithms of ActStudio software version 2.1.2 (Condor Instruments, SP, Brazil), which included automatic detection of sleep-wake cycles and off-wrist periods, parameter calculations, and export of study actograms. Sleep-wake cycles were visually inspected by blinded investigators to confirm the algorithm determinations, with no data altered or manipulated.

Circadian Function Index (CFI) was used as an overall measure of circadian robustness. This index incorporates three important parameters: interdaily stability (IS), which reflects the consistency of rhythms over multiple days and their synchronization with the light-dark cycle (ranging from 0 to 1 with higher values indicating greater stability); intradaily variability (IV), which measures the fragmentation of circadian rhythms (ranging from 0 to 2 with higher values indicating greater fragmentation); and relative amplitude (RA), the difference between the average activity during the most active 10 consecutive hours (M10) and the least active 5 consecutive hours (L5).^40^ Off-wrist values were not included in the analysis (ActStudio software; calculations based on Witting *et al.*^41^). CFI represents the average of IS, RA, and IV (inverted and normalized between 0 and 1) and ranges from 0 (absence of rhythmicity) to 1 (robust circadian rhythm). To calculate CFI, the software uses IV and IS based on data sampled every 60 minutes (IV60 and IS60) and the average day RA (ActStudio software; calculations based on Ortiz-Tudela *et al.*^40^). When assessing IV and IS parameters individually, we calculated their mean using all sampling intervals (ISm and IVm). This method has been reported by Gonçalves *et al.*^42^ to more precisely detect sleep-wake cycle fragmentation and synchronization.

A cosine curve fitting was applied to actigraphy data to extract parametric measures such as amplitude (difference between the peak and the average function value) and acrophase (time of peak activity).^43^ Additionally, sleep efficiency (percentage of time spent asleep while in bed) and total sleep time (overall duration of sleep) were also automatically calculated (ActStudio software version 2.1.2).

### Animal handling

The mouse model of MJD carrying the full human *ATXN3* gene with 84 CAG repetitions (YAC-MJD; MJD YAC84), with two copies (hemizygous; MJD YAC84.2; Q84.2/-) or four copies (homozygous; MJD YAC84.2/84.2; Q84.2/Q84.2) of the transgene, was used to conduct *in vivo* studies.^32^ Control C57BL/6 mice were acquired from Charles River Laboratories and MJD YAC84 mice were acquired from the Jackson Laboratory. A colony of transgenic mice was established in the licensed animal facility (International Animal Welfare Assurance number 520.000.000.2006) at the Center for Neuroscience and Cell Biology of the University of Coimbra. Animals were housed in a temperature- (22 ± 2 °C) and humidity- (55 ± 15 %) controlled room, on a 12 h light-dark (LD) cycle. Food and water were provided *ad libitum*. This study included male and female mice, aged 5 to 13 months. The experiments were carried out in accordance with the European Union Community directive (2010/63/EU) for the care and use of laboratory animals. Researchers received adequate training (Federation of European Laboratory Animal Science Associations (FELASA)-certified course) and certification from Portuguese authorities responsible for the regulation of animal experimentation (Direcção Geral de Alimentação e Veterinária) to perform the experiments. This study is part of a research project that was approved by the Center for Neuroscience and Cell Biology ethics committee (ORBEA_289_2021/10122021) and DAGV (DGAV 0421/000/000/2022).

### Locomotor activity assays

Wheel-running activity was assessed using a custom-made mouse cage, fitted with a running wheel containing a small neodymium magnet. Once per revolution, the magnet triggered a reed switch placed outside the cage allowing data to be collected via an Arduino interface running custom code.

After 4 days of handling and habituation, mice were individually placed in wheel-running cages on a 12 h LD cycle (6 h–18 h/18 h–6 h). Following 3 days of habituation to the wheel-running cage, the number of wheel turns was recorded for an additional 7 days to establish baseline activity. After this period, the light cycle changed to either: i) a 12:12 h dark/dark (DD) free-running condition (“DD” protocol); or ii) a new 12 h LD cycle, after an abrupt 4 h phase advance in light entrainment (“jet-lag” protocol), as described elsewhere.^31^ Raw data containing wheel-running activity was pre-processed by a custom-made python script and actograms were generated and analysed in specific time periods using the ImageJ plugin ActogramJ.

In the DD protocol, the inactive period of the mice was considered from 7 h to 14 h. Total day activity was also analysed for the entire study and further divided into two periods: from 00 h to 12 h and from 12 h to 24 h. Fragmentation was analysed from 18 h to 24 h during the days of the LD cycle and defined as the number of bouts (intervals of 18 consecutive minutes) in which wheel-running revolutions were inferior to 90 (equivalent to a threshold of 5 laps per minute), adapted from pre-established criteria for ClockLab (Actimetrics, Wilmette, IL).^44^ Imprecision (variability in the observed onset of activity compared to the linear expected onset) and innate circadian period calculation (duration of one full cycle of activity, from the start of one activity phase to the start of the next), were calculated based on daily activity onset during the free-running period in the DD protocol. A regression line was created using the activity onset data from each animal, which served as the mathematical foundation for this analysis. Activity onset was calculated in ActogramJ after a Gaussian distribution smoothing of the data, with the median of all non-negative activity values used as the threshold. With the slope of the regression line being used to calculate the period, and the r square to calculate imprecision, this method allows for quantification of common circadian rhythm markers without the use of subjective eye-fitted lines or commercially available software.

The number of days needed by each animal to adjust to the 4 h advance in light entrainment in the jet-lag protocol was determined by investigators blinded to the experimental groups.

### Telemetric temperature probes implantation

Wild-type (WT) and homozygous male animals weighing more than 22 g were selected for the implantation of electronic telemetric capsules (Anipill; BodyCap, Paris, France). The sterile pills were activated according to the manufacturer’s instructions and implantation was carried out by adopting a standard operating procedure provided by the manufacturer and the available published protocols.^45,46^ Approximately 30 minutes before surgery, each mouse was injected with buprenorphine as pre-operative analgesia. Mice were anaesthetized individually with isoflurane (induction chamber, 5 % in 2 L/min oxygen; Abbott) until loss of the righting reflex. Mice were then placed in a dorsal position on a heating pad and anaesthesia maintained via nose cone (1.5-2 % in 2 L/min oxygen). The fur in the abdominal area was shaved, and the skin disinfected with chlorhexidine and 70 % alcohol. A sagittal midline incision of 1 to 1.5 cm along the *linea alba* was performed and the capsule was gently positioned in the peritoneal cavity, on the opposite side of the cecum. After insertion, muscle (synthetic absorbable monofilament suture, Monosyn 6-0) and skin (natural non-absorbable multifilament, Silkam 5-0) were sewed separately. To avoid fighting after surgery and during the experimental period, animals were returned to an individual clean cage and were allowed to recover with the help of heating pads and moist food pellets. Post-operative analgesia was administered for 3 consecutive days after surgery: buprenorphine was given every 8 hours for the first 48 hours, followed by a single dose of carprofen after 48 hours. CBT was recorded wirelessly every 5 minutes for 1 month in animals housed in a room maintained at 23 °C. Only data from the last 16 days were analysed. CBT values were compared between groups and CircaCompare was used to estimate circadian parameters.^47^ This novel statistical method not only allows the appropriate fitting of recent cosinor regression-based methods but also facilitates the comparison of circadian parameters between the two groups.^48,49^

### Tissue preparation and immunofluorescence

Mice from the DD activity study were housed in complete darkness for 24 h prior to whole-brain sample collection.^31^ Mice were sacrificed with a lethal intraperitoneal administration of ketamine and xylazine, followed by intracardiac perfusion with ice-cold PBS and fixation with 4 % paraformaldehyde (Sigma). Whole-brain samples were quickly harvested, post-fixed in 4 % paraformaldehyde for 24 h, and then transferred to a 25 % sucrose/PBS solution for 36-48 h for cryoprotection and dehydration. Brain samples were then frozen at −80 °C. Coronal sections (25 µm) were cut using a cryostat (Leica CM3050S, Leica Microsystems) at −20 °C. Sections were collected in anatomical series and were stored in 48-well plates in PBS supplemented with 0.05 % (m/v) sodium azide, at 4 °C, until further processing.

Free-floating immunofluorescence was initiated by washing the selected brain sections with PBS and then incubation for 1 h in a blocking and permeabilizing solution of 3 % BSA/10 % goat serum/0.3 % triton X-100 in PBS at room temperature. Sections were then incubated overnight at 4 °C with two primary antibodies: rabbit polyclonal anti-arginine vasopressin (AVP; 1:800, #20069, Immunostar) and mouse monoclonal anti-vasoactive intestinal peptide (VIP; 1:250, sc-25347, Santa Cruz Biotechnology) diluted in blocking solution. Sections were washed in PBS and incubated for 2 h at room temperature with the secondary antibodies coupled to fluorophores (Alexa Fluor 568 conjugated goat anti-rabbit and Alexa Fluor 488 conjugated goat anti-mouse, 1:200) diluted in blocking solution. Nuclei counterstaining was performed with Hoechst 33342 (2 μg/mL; Invitrogen Molecular Probes) during the described secondary antibody incubation. Sections were washed with PBS, mounted on gelatine-coated slides and coverslipped with mounting medium (Dako). Images were acquired using an Axio Scan.Z1 equipped with a Plan-Apochromat 20×/0.8 M27 objective. Analysis was performed with QuPath Software. The regions of interest (ROI) of SCN and paraventricular nucleus (PVN) were defined by dense Hoechst staining and performed according to The Paxino’s Mouse Brain Atlas.^50^ For each mouse, VIP and AVP integrated densities of at least 14 unilateral SCN and 12 unilateral PVN were measured (product of area and mean fluorescence value; arbitrary units, all values divided by 10^8^; ROI: 0.50 µm per pixel). In the same ROI, AVP cell counting was determined using the “Positive cell detection” tool of QuPath. Cells were detected using Hoechst nuclear labelling (threshold of 1500) and positive AVP cells were automatically counted when bright marking was present (single threshold for cell AVP mean: 7000 for PVN and 9000 for SCN; thresholds defined by two independent researchers). To ensure analysis of similar SCN positions between groups, only ROI of 500 to 900 cells, without autofluorescence contaminating the ROI, were included in the final comparison. Integrated density values and percentage of positive cells of all the unilateral regions (left and right) were averaged for each region (SCN and PVN) of each animal. Quantifications were performed by a researcher blinded to the experimental groups.

### Cell culture

#### Neuro2a cell culture and transfection

A mouse neuroblastoma (neural crest-derived) cell line (N2a cell line; American Type Culture Collection) was maintained in Dulbecco’s modified Eagle’s medium (DMEM; Sigma-Aldrich) supplemented with 10 % heat-inactivated foetal bovine serum (FBS; Life Technologies) and 1 % penicillin-streptomycin (LifeTechnologies), at 37 °C under a humidified atmosphere containing 5 % CO_2_. For cell transfection, 4×10^4^ cells were plated on 96-well cell culture multiwell plates coated with poly-D-lysine (Sigma-Aldrich). After 24 hours, cells were transfected with a mixture of DNA/polyethyleneimine (PEI) complexes (MW40000, PolySciences) containing *Bmal1*-Luc reporter; *Per2*-Luc reporter; WT ATXN3 (LV-PGK-ATXN3 27Q); or mutATXN3 (LV-PGK-ATXN3 72Q) plasmids.^31,51^ Complex formation was performed by combining 30 ng of reporter plasmids with 60 ng of WT ATXN3 or mutATXN3 constructs in 20 µL of serum-free DMEM and 1 µL of PEI (1 mg/mL) per well, as adapted from Chang & Guarente^31^. DNA/PEI mixtures were vortexed for 10 s and incubated at room temperature for 10 min. Afterwards, DNA/PEI complexes were diluted in serum-free DMEM and added to the cells in culture.

#### Bioluminescence assays

After transfection (4 h), cells were synchronized with 50 % horse serum for 2 h followed by medium replacement with DMEM with low-FBS (1 %) for 24 hours before luciferase assays, as previously described.^31^ Cells were washed with PBS and incubated with Bright-Glo^®^ reagent (Promega) diluted in PBS (1:2). To ensure cell lysis, samples were incubated with the reagent for at least 3 min before reading on FLUOstar Omega Microplate Reader (BMG LABTECH). At least 3 reads, every 3 minutes, were performed on each sample, and the average values were considered for analysis. Each sample was present in triplicate in each analysis and the assays were repeated 4 different times.

### Gene expression analysis

#### Tissue collection and RNA extraction

Mice were anaesthetized with isoflurane (Abbott) and decapitated at specified Zeitgeber Times (ZT, 3 h, 11 h, and 19 h). Red dim light was used during the night timepoint. Brains were quickly harvested and the cerebellum and hypothalamus were dissected, snap-frozen, and stored at −80 °C until processing. Tissue homogenisation was performed with TRIzol™ Reagent (Invitrogen) and chloroform (Merck), and RNA was isolated with NucleoSpin RNA II isolation kit (Machery Nagel) according to the manufacturer’s instructions. RNA purity and concentration were assessed with a Nanodrop 2000 Spectrophotometer (Thermo Fisher Scientific). RNA samples were stored at −80 °C until further use.

#### cDNA synthesis and quantitative real-time PCR

RNA samples were converted into cDNA, using the iScript cDNA Synthesis Kit (Bio-Rad), according to the manufacturer’s instructions, and stored at −20 °C. The mRNA levels of core clock genes, *Bmal1*, *Per2*, *Clock*, and *Cry1* were assessed by real-time quantitative reverse transcriptase polymerase chain reaction (qRT-PCR), using the CFX96 Touch Real-Time PCR Detection System (Bio-Rad). Reactions of 10 µL were prepared using 4 μL of template cDNA, 0.5 μL of 10 µM forward and reverse primers (final concentration of 0.5 µM; Supplementary Table 2), and 5 μL of the SsoAdvanced SYBR Green supermix (Bio-Rad). Reactions were performed according to the manufacturer’s recommendations: 95 °C for 30 seconds, followed by 45 cycles at 95 °C for 5 seconds, and 56 °C to 60 °C (annealing temperatures specified in Supplementary Table 2) for 15 seconds. The melting curve protocol started immediately after amplification, with slow heating, starting at 65 °C with an increment of 0.5 °C/5 s up to 95 °C. Oligonucleotide sequences for the amplification of clock genes were as we previously reported.^52^ Each assay included a non-template control (NTC), a no reverse transcription (NRT) control, and a standard curve for each target gene. Amplification efficiencies and threshold cycles (CT) were automatically determined by Bio-Rad CFX Maestro software. Relative mRNA quantification was performed using the ΔCT method.^53^ The mRNA expression data is shown as ΔΔCT values, relative to the average ΔCT of all time points of all control samples, as previously described.^54^ Data normalization was performed using the *Hprt* housekeeping gene.

### Statistical analysis

Results are expressed as mean ± standard error of the mean (SEM). Statistical analysis was performed using GraphPad Prism 8 (version 8.0.2, GraphPad). Data distribution was assessed using the Shapiro–Wilk test and the quartile–quartile plot (QQ-plot) was visually inspected. To evaluate correlations, Pearson correlations were used to determine the Pearson correlation coefficient (r), goodness of fit (R square), and *P* value. Paired or unpaired t tests were performed for comparisons between two groups, while one-way ANOVA followed by Dunnett’s post hoc test was used to compare multiple conditions. *P*-values are reported. In the analysis of CBT, data was analysed using multiple t tests, corrected for multiple comparisons through the Holm-Sidak method, and the adjusted *P* value was reported. For the analysis of the circadian parameters of CBT, we used CircaCompare ShinyR application (available here: https://rwparsons.shinyapps.io/circacompare/).^47^ All statistical tests performed were two-tailed, with the statistical threshold set at \**P* < 0.05, \*\**P* < 0.01, \*\*\**P* < 0.001, and \*\*\*\**P* < 0.0001. No values were excluded. Specific statistical parameters are detailed in the figure legends.

## Results

### Disease progression in patients with MJD is associated with a circadian rhythm decline

We analysed the integrity of circadian rhythms in MJD patients at different disease stages by evaluating their rest-activity cycles using an actigraphy device over a 2-week period (Fig. 1A). The MJD patients were characterized in what concerns sex, age, number of CAG repeats, disease duration, clinical condition (SARA score, INAS), functional capacity (ADL, CCFS) and sleep quality (PSQI). This study included five females and two males, aged 53 to 70 years (age not correlated with disease severity), with 63 to 68 CAG repeats, and a disease duration of 7 to 18 years (Fig. 1B). According to the SARA score, patients were classified as mildly ataxic (dark blue), moderately ataxic (purple), or severely ataxic (red; Fig. 1B).

**Figure 1.**
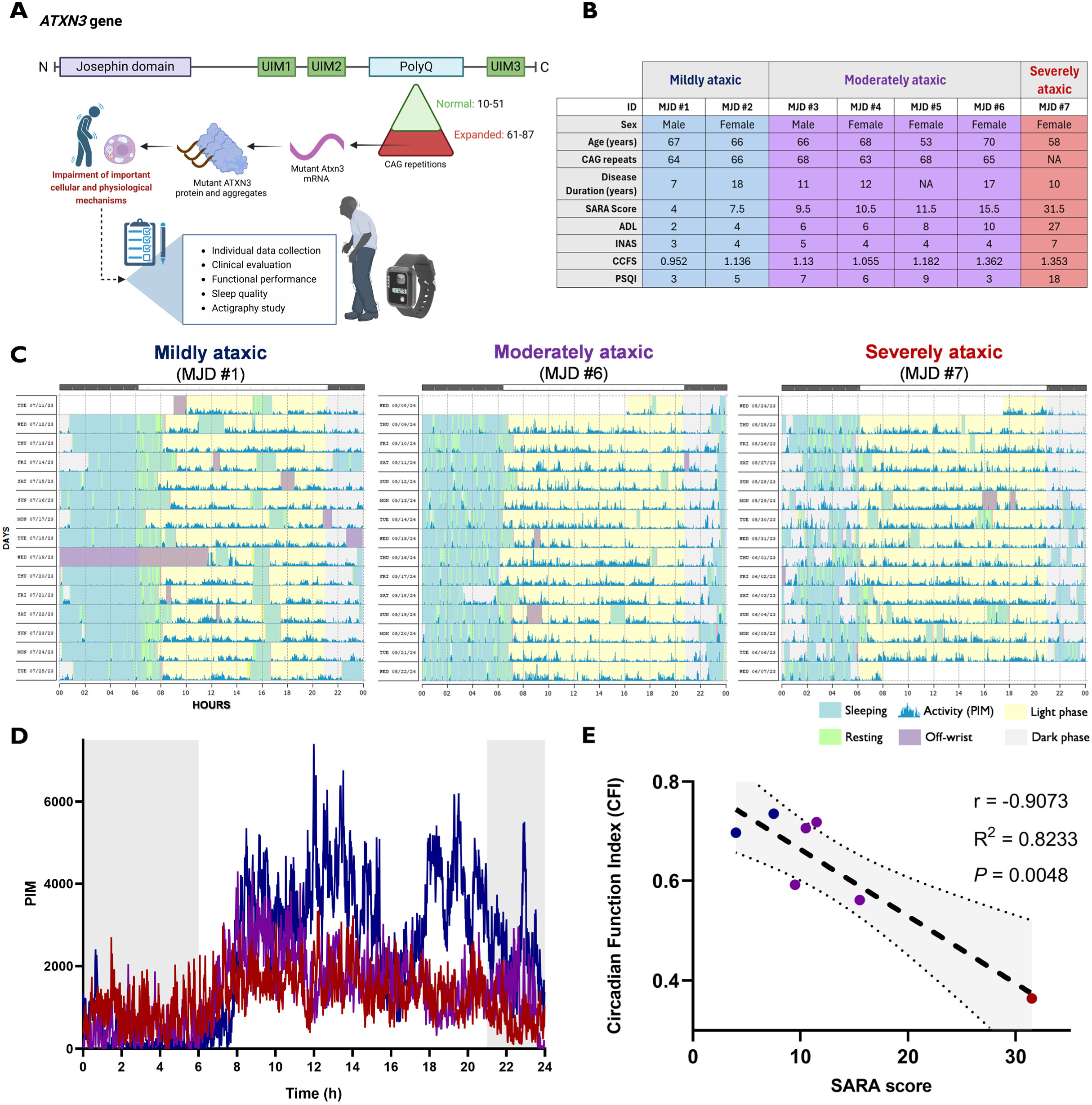
Disease progression in patients with MJD is associated with a circadian rhythm decline. (A) Schematic representation of MJD pathological features development and actigraphy study design. (B) Characteristics of MJD patients enrolled in the study, including age, sex, number of CAG repetitions, disease duration, and clinical scales score (SARA, ADL, INAS, CCFS, and PSQI). Participants were ordered and stratified by SARA score as mildly ataxic (3.0 ≤ SARA ≤ 7.5; dark blue), moderately ataxic (8.0 ≤ SARA ≤ 23.5; purple) or severely ataxic (SARA ≥ 24.0; red). (C) Representative actograms of 3 MJD patients in different disease states followed for two weeks. Mild ataxia represented by MJD #1 patient, moderate ataxia by MJD #6 patient, and severe ataxia by MJD #7 patient. Blue peaks represent activity (PIM) normalized for each patient, light blue sleeping states, light green resting states, light yellow light phases, light grey dark phases, and purple off-wrist periods. Black (night) and white (day) bars are defined according to sunrise and sunset times of the study. (D) Representative average daily graphs depicting the hours of higher and lower activity across the different disease states. Mean PIM value calculated for each minute based on the raw data. (E) Correlation of SARA score with circadian function index (CFI), reflecting the overall decline in circadian robustness along the disease course. Data distribution assessed using the Shapiro– Wilk test. Pearson correlation was used to determine the correlation coefficient (r), goodness of fit (R square), and *P* value. 95 % confidence intervals are shown. Abbreviatures: Activities of Daily Living (ADL); Composite Cerebellar Functional Severity score (CCFS); Inventory of Non-Ataxic Signs (INAS); Pittsburgh Sleep Quality Index (PSQI); Scale for the Assessment and Rating of Ataxia (SARA).

Representative actograms of MJD patients stratified in the three different disease states suggested a progressive disruption of the rest-activity rhythms within the course of the pathology (Fig. 1C). This hypothesis was reinforced when inspecting the respective average daily graphs of activity (PIM) over time (Fig. 1D). To investigate this potential progressive circadian decline, we evaluated the Circadian Function Index (CFI) of each MJD patient. CFI is a sensitive and accurate integrated measure of the overall circadian robustness and represents the normalized average of the intradaily variability (IV60; fragmentation of circadian rhythms), interdaily stability (IS60; rhythm consistency and synchronization across the days) and relative amplitude (RA; difference between mean activity during the most active consecutive 10 h, M10, and the least active 5 consecutive hours, L5).^40,55^ We observed a progressive decline in the robustness of rest-activity rhythms with disease progression in MJD patients, as evidenced by a strong negative correlation between the CFI and the SARA score (r = −0.9073, *P* = 0.0048; Fig. 1E). Similar negative correlations were found with ADL and INAS scores (ADL: r = −0.9102, *P* = 0.0044; INAS: r = −0.8751, *P* = 0.0099; Supplementary Fig. 1A). Additionally, trends were observed with functional performance (CCFS: r = −0.7255, *P* = 0.0650) and sleep quality (PSQI: r = −0.7301, *P* = 0.0624; Supplementary Fig. 1A). When looking at the individual nonparametric parameters, ISm was decreased with MJD progression (r = −0.8421, *P* = 0.0175), while IVm followed the opposite profile (r = 0.8004, *P* = 0.0306; Supplementary Fig. 1B). Importantly, although M10 suggested that daytime activity levels are not strongly correlated with the progression of MJD, the L5 correlation indicated that activity is increased during the rest phase despite aggravated ataxia symptoms (r = 0.7729, *P* = 0.0416; Supplementary Fig. 1B). This is in agreement with the activity profile observed during the night in the representative graphs of MJD patients at later stages of the disease (Fig. 1C-D).

Amplitude (difference between the peak and the average function value) and acrophase (time of peak activity) were calculated using cosine curve fitting of the activity data.^43^ MJD patients showed a trend towards decreased amplitude and a phase advance, indicating an earlier peak activity time as the disease progressed (Supplementary Fig. 1C). Furthermore, sleep efficiency and total sleep time diminished with the increase in SARA score (sleep efficiency: r = −0.8930, *P* = 0.0068; total sleep time: r = −0.8493, *P* = 0.0156; Supplementary Fig. 1D). No alterations in the number of awakenings were observed (data not shown).

These results suggest that the clinical progression of MJD is associated with a pronounced decline in circadian function.

### The YAC-MJD transgenic mouse model shows impaired circadian activity rhythms and decreased re-entrainment capacity

Following the observation that patients with MJD show a progressive circadian decline along the disease course, we investigated the neurobiological mechanisms behind this dysfunction in a mouse model of MJD — a YAC-MJD transgenic mouse model, which encodes the full-length mut*ATXN3* gene under the control of its own promoter — by conducting wheel-running experiments under different light-dark protocols (Fig. 2A).

**Figure 2.**
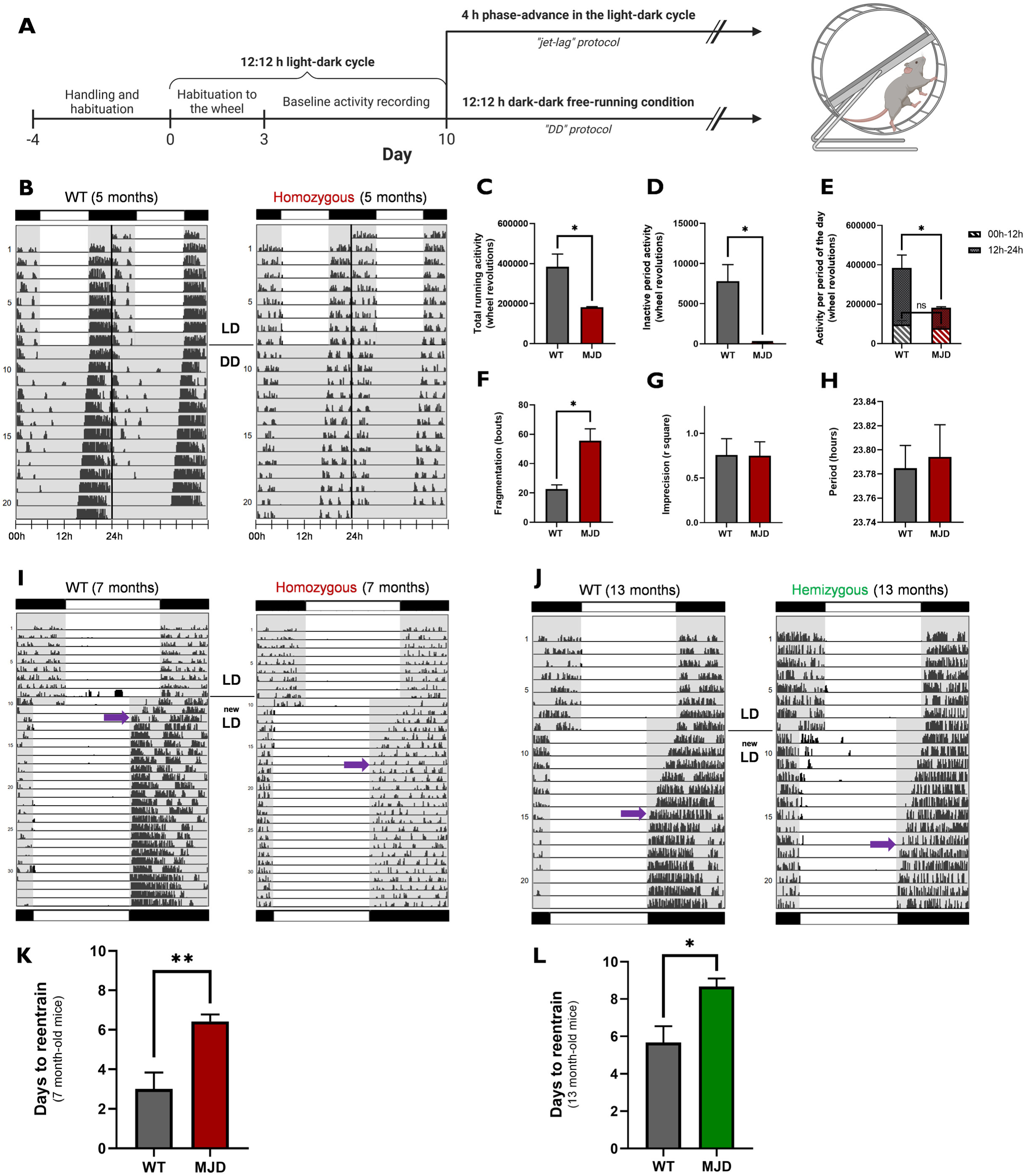
YAC-MJD transgenic mouse model shows impaired circadian activity rhythms and decreased re-entrainment capacity. (A) Schematic representation of the protocol for wheel-running experiments performed with the YAC-MJD transgenic mouse model under different light-dark conditions. (B) Representative double-plotted actograms of wheel-running activity of 5-month-old male MJD YAC84.2/84.2 (homozygous, *n* = 3) and wild-type control mice (WT, *n* = 3) in constant darkness, after 8 days of light-dark (LD) entrainment. Total wheel revolutions determined (C) during the complete study, (D) in 7 hours of the inactive period, (E) and for the two parts of the day. (F) Number of bouts, intervals of 18 consecutive minutes, where wheel-running revolutions were inferior to 90. Fragmentation was determined in 6 representative hours of the active phase during the days of LD entrainment. Homozygous mice showed (F) increased fragmentation and (C-E) significantly reduced levels of activity in specific parts of the day. During the free-running period, under the dark-dark (DD) light scheme, the regression line of the activity on-set was used to determine (G) imprecision, r square representing variability in the start of activity, and (H) innate circadian periods of the animals. (I, J) Actograms from mice subjected to a 4-hour phase advance jet lag experiment. Representative results of 7-month-old male homozygous (*n* = 6) compared to WT control mice (*n* = 5) are shown on the left panel (I) while 13-month-old male MJD YAC84.2 (hemizygous, *n* = 3) and WT control mice (*n* = 3) are shown on the right (J). (K) Homozygous mice required 3.42 additional days to re-entrain to the new LD cycle, and (L) Hemizygous mice required 3.00 more days for the same re-entrainment, compared to their respective WT control mice. Purple arrows represent the day of re-entrainment. When the arrow is present between two days it is the result of the average quantification of two independent researchers. Data in (C-H) and (K, L) is presented as mean ± SEM. Grey rectangles represent the dark phase. Unpaired t tests were performed for comparison between groups. Statistical significance is shown as \**P* < 0.05, \*\**P* < 0.01. ns: not significant.

In the first experiment, 5-month-old mice were entrained on a 12 h LD cycle for the first 8 days and then transferred into a DD environment for 2 weeks (Fig. 2B; Supplementary Fig. 2A). As expected, homozygous mice showed a 53 % reduction in overall activity (total wheel revolutions) throughout the study, when compared to WT mice (Fig. 2C). This reduction was also evident over three representative days under the two different light schemes conducted (Supplementary Fig. 2B). Furthermore, homozygous mice also showed a drastic decrease in activity during 7 h of the inactive period throughout the entire study (Fig. 2D). Interestingly, no significant differences in activity were observed during the late-night period (Fig. 2E; Supplementary Fig. 2A). Importantly, we observed a fragmentation 2.46 times higher in the activity of homozygous mice, compared to WT mice (Fig. 2F; Supplementary Fig. 2A). A regression analysis of the activity during the free-running period revealed no alterations in imprecision (Fig. 2G) or innate circadian periods (Fig. 2H) of homozygous mice, compared to WT mice. Throughout the study, as previously described, homozygous mice showed loss of body weight gain, compared to WT mice; yet neither the homozygous nor WT mice lost body weight after the activity evaluation (Supplementary Fig. 2C).^26^

To further characterize the circadian phenotype of the YAC-MJD transgenic mouse model, 7-month-old homozygous (Fig. 2I), 13-month-old male hemizygous (Fig. 2J), and their respective WT controls, were submitted to a well-established jet lag experiment in which light entrainment was abruptly advanced by 4 h, as described by others.^31,56,57^ We found that homozygous mice required 3.42 additional days for re-entrainment to the new LD cycle compared to WT mice (Fig. 2K). Importantly, this represents double the time needed by the WT controls (7-month-old mice; WT: 3.00 ± 0.84 days; homozygous: 6.42 ± 0.35 days; Fig. 2K). On the other hand, hemizygous mice required 3.00 more days for re-entrainment compared to their aged-matched controls (Fig. 2L). This increase was less pronounced compared to the one observed in homozygous mice, but these older animals showed overall longer re-entrainment times (13-month-old mice; WT: 5.67 ± 0.88 days; hemizygous: 8.67 ± 0.44 days; Fig. 2L).

These findings show that circadian activity rhythms and re-entrainment capacity are compromised in the YAC-MJD transgenic mouse model, suggesting a central circadian dysfunction.

### YAC-MJD homozygous mice show a disruption in core body temperature rhythms

After observing that activity rhythms are disrupted in the YAC-MJD transgenic mouse model, we sought to determine whether the biological output of circadian rhythms, specifically core body temperature (CBT), is also affected.

For that purpose, we surgically implanted telemetric temperature data loggers into the abdomen of 7-month-old homozygous and WT mice for continuous recording of CBT in a freely-moving environment (Fig. 3A). CBT was recorded every 5 minutes and analysed over the last 16 days of the experiment (Fig. 3B). Mean CBT values were calculated for every timepoint and individual values for each mouse were plotted in an average day graph of both genotypes (Fig. 3C). Strikingly, homozygous mice showed an increase in CBT at the beginning of their active phase, between 20 h and 22 h, with a maximum increase of 1.08 °C and adjusted *P* values as significant as *P* = 0.000002 (Fig. 3C; Supplementary Table S1). Moreover, median CBT was also increased in the homozygous compared to WT mice (Fig. 3D). We further compared the circadian parameters of CBT rhythms over the entire 384-hour recording period, as described by Parsons *et al*.^47^ Both homozygous and WT mice displayed rhythmic CBT profiles (Table 1). As displayed in the raw data comparison, homozygous mice showed a higher Midline Statistic Of Rhythm (MESOR; representing the rhythm-adjusted mean temperature), and increased amplitude estimates (Table 1). Additionally, homozygous mice showed a phase advance of more than 1 h in the time at which CBT peaks (acrophase) compared to WT controls (WT: 3.26 h; homozygous: 1.95 h; Table 1). Differences in circadian parameters, in particular acrophase and amplitude, were visible when fitting the CBT values for the average day using CircaCompare (Supplementary Figure 3A).

**Figure 3.**
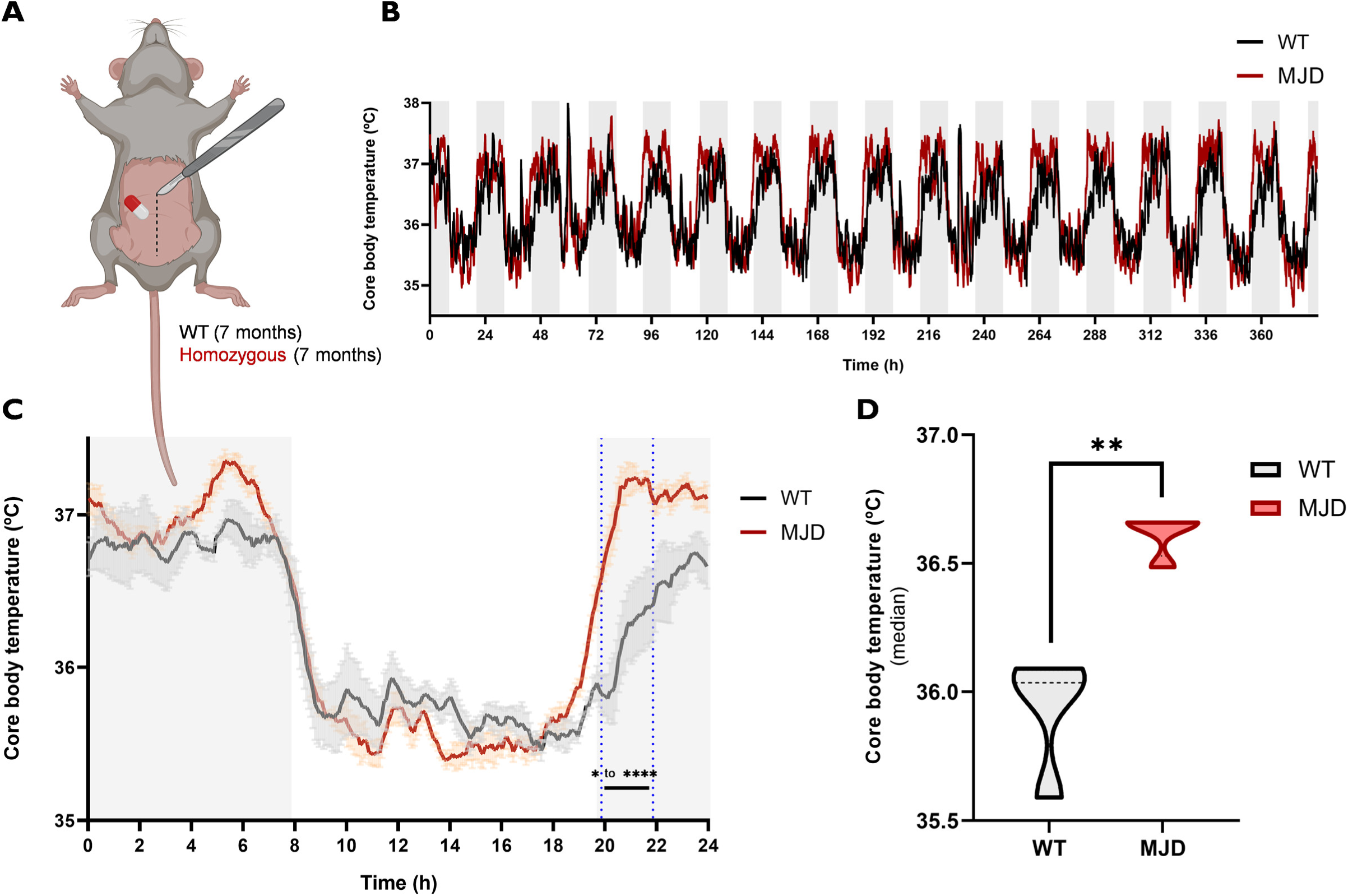
YAC-MJD homozygous mice show a disruption in core body temperature (CBT) rhythms. (A) Representative scheme of a surgical abdominal implementation of a temperature data logger for CBT monitoring. (B) Graphical representation of CBT recorded every 5 minutes for 16 days (*n* = 4). Lines represent the mean of 7-month-old male MJD YAC84.2/84.2 (homozygous) and wild-type (WT) control mice. (C) Plot of average CBT values along the day. Homozygous mice showed a very evident increase in temperature at the beginning of the active phase. (D) Violin plot of median CBT showing the overall increase of CBT in the homozygous mice compared to WT. Grey rectangles indicate the dark phases. Multiple t tests were performed for comparison of CBT values between groups at each time point (C), corrected for multiple comparisons using the Holm-Sidak method. CBT median values (D) were compared between groups by an unpaired t test. Statistical significance was set as: \**P* < 0.05, \*\**P* < 0.01, dashed blue lines (* to ****): \**P* < 0.05 to \*\*\*\**P* < 0.0001.

**Table 1.**
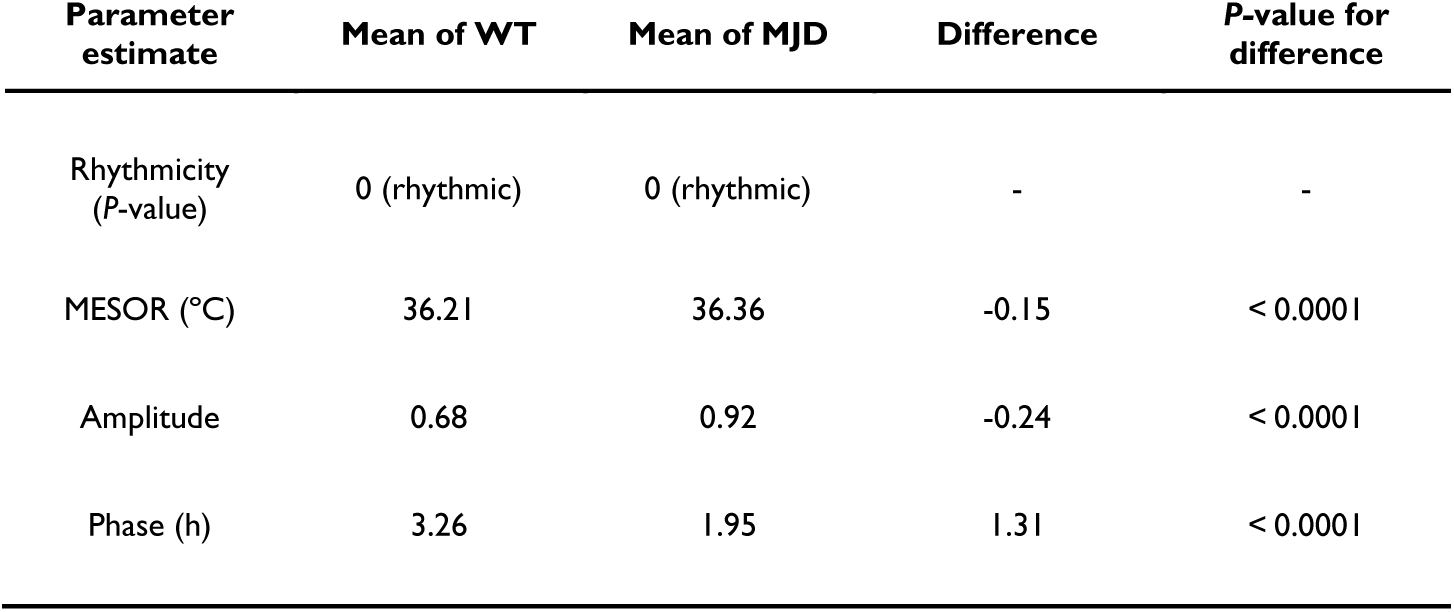
Circadian rhythm parameters of core body temperature data of the YAC-MJD homozygous mice determined using CircaCompare for the total 16 days.

Altogether, these results suggest that CBT regulation is robustly and significantly affected in the YAC-MJD homozygous mice.

### YAC-MJD homozygous mice show reduced levels of the synchronizing neuropeptides VIP and AVP in the hypothalamic SCN and PVN

Given that circadian rhythms of activity and CBT are significantly affected in MJD, we hypothesized that alterations in the central regulator, the SCN, are key contributors to these impairments.

For that, we evaluated AVP and VIP levels in the SCN and PVN of the 5-month-old mice subjected to the DD wheel-running activity study. AVP and VIP are crucial to regulate the circadian rhythms of several physiological functions, including rest-activity cycles and CBT.^58–60^ Their dysfunction in the master clock has been suggested as one of the main mechanisms behind circadian disturbances in neurodegenerative disorders.^22,61^ Fluorescent immunohistochemical analysis of VIP and AVP demonstrated strong levels of VIP and AVP in the SCN, PVN, and supraoptic nucleus of the hypothalamus (Fig. 4A), including the SCN-PVN projections, through the subparaventricular zone, necessary to regulate the daily rhythms of endocrine and autonomic functions, including temperature regulation (Fig. 4B).^62–65^ The SCN of homozygous mice showed strongly reduced levels of VIP (WT: 4.99 ± 0.20 AU; homozygous: 4.05 ± 0.52 AU; Fig. 4C) and AVP (WT: 6.23 ± 0.24 AU; homozygous: 5.20 ± 0.34 AU; Fig. 4D) immunoreactivity compared to WT mice (Fig. 4E). The AVP decrease in the SCN corresponded to a 65 % reduction in AVP positive cells (WT: 28.25 ± 3.62 %, equivalent to mean 2247 positive cells per mm^2^; homozygous: 9.86 ± 3.13 %, equivalent to mean 753 positive cells per mm^2^; Fig. 4F). The percentage of AVP positive cells and cell counts in WT mice were similar to previously reported values.^66^ In another relevant hypothalamic area, the PVN (important for the regulation of circadian rhythms and synchronization across the peripheral clocks^67^), the reduction of AVP levels was even more pronounced in the homozygous mice compared to WT mice (Fig. 4G-H). The AVP decrease in the PVN corresponded to a dramatic 78 % reduction in AVP positive cells (WT: 12.11 ± 1.15 %, equivalent to mean 884 positive cells per mm^2^; homozygous: 2.66 ± 1.01 %, equivalent to mean 183 positive cells per mm^2^; Fig. 4I). As expected, VIP immunofluorescence was minimal in the PVN, and no differences were observed between the two experimental groups (data not shown).

**Figure 4.**
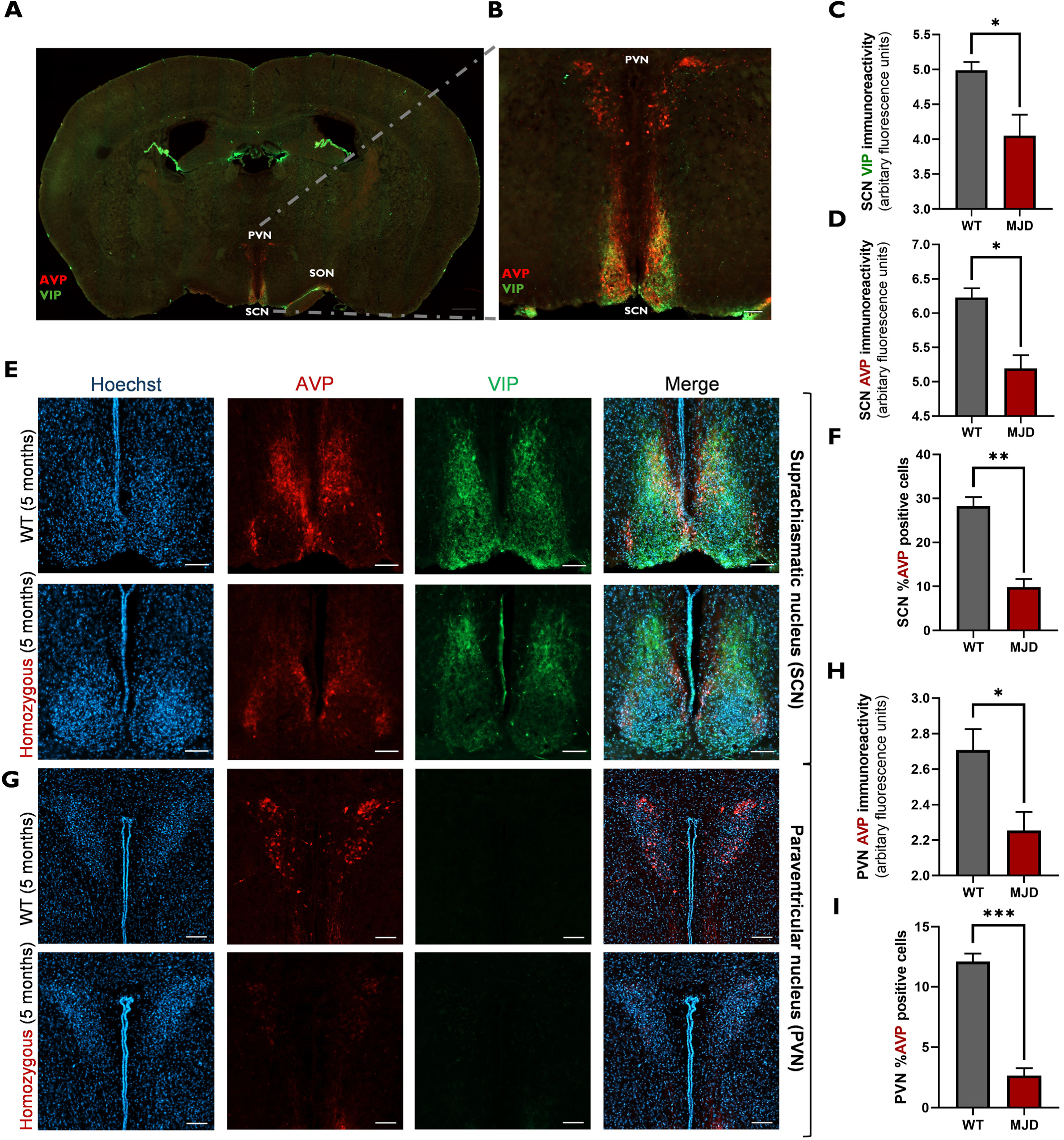
YAC-MJD homozygous mice show reduced levels of the synchronizing neuropeptides vasoactive intestinal polypeptide (VIP) and arginine vasopressin (AVP) in the hypothalamic suprachiasmatic nucleus (SCN) and paraventricular nucleus (PVN). (A) Fluorescent immunohistochemical analysis of AVP and VIP illustrating the levels of these neuropeptides in the SCN, PVN, and the supraoptic nucleus (SON) of the hypothalamus. (B) SCN AVP and VIP neuronal projections to the PVN, essential to regulate circadian rhythms. Representative co-immunofluorescence images of AVP (in red) and VIP (in green) in the (E) SCN and (G) PVN of MJD YAC84.2/84.2 (homozygous) and wild-type (WT) mice upon constant darkness. (C) Quantification of VIP and (D) AVP immunoreactivity showing decreased levels of both neuropeptides in the SCN and (H) of AVP in the PVN of homozygous mice compared to control animals. (F, I) A 65 % and 78 % reduction of AVP positive cells was observed in the SCN and PVN, respectively, of homozygous mice compared to WT control mice. Data in (C, D, F, H, I) is presented as mean ± SEM. Values between groups were compared by unpaired t tests (*n* = 3 animals per group; >12 PVN and >14 SCN unilateral regions, analysed for each animal). Statistical significance was set as: \**P* < 0.05, \*\**P* < 0.01, \*\*\**P* < 0.001. Scale bars represent (A) 500 μm, (B) 100 μm, (E, G) 50 μm.

In conclusion, we found evidence of a drastic decline in the neuropeptides regulating the central clocks, particularly in the SCN and PVN, which may underlie the circadian dysfunction observed in the YAC-MJD transgenic mouse model.

### YAC-MJD homozygous mice show a decline in the expression of core clock genes in the cerebellum

The master clock and its neuropeptides are essential for the coordination of peripheral clocks throughout the brain and body via cell-autonomous transcriptional/translational negative feedback loops of clock gene expression.^64^ Given that YAC-MJD mice exhibited decreased levels of the synchronizing neuropeptides VIP and AVP in the master clock, we hypothesized that clock gene expression is also impaired, contributing to the circadian rhythm alterations in MJD.

To test this hypothesis, we evaluated the expression levels of four core clock genes (*Bmal1*, *Clock*, *Per2*, *Cry1*) at ZT 3 h, ZT 11 h, and ZT 19 h in two regions of the brain: in the hypothalamus, which houses the master clock; and the cerebellum – one of the most affected brain regions in MJD and one of the areas of the brain with more pronounced rhythms in gene expression.^5,9,68^ Despite the heterogeneity of the experimental groups, we observed alterations in clock gene expression in the 10-12 month-old hemizygous mice compared to WT controls (Supplementary Figure 4A-B). A significant increase in the night timepoint (ZT 19 h) was observed for *Clock* expression in both the hypothalamus (Supplementary Figure 4A) and cerebellum (Supplementary Figure 4B) of hemizygous mice compared to WT controls. Moreover, an increased number of tendencies for dysregulation (*P* < 0.1) were observed in the cerebellum (*Bmal1* ZT 19 h; *Per2* ZT 19 h; *Cry1* ZT 19h; Supplementary Figure 4B) compared to the hypothalamus (*Per2* ZT 19 h; Supplementary Figure 4A). Nevertheless, we observed considerable variability within the study population, as evident by the data dispersion of the mRNA expression graphs.

To further explore this hypothesis, we analysed the expression of these core clock genes in the cerebellum — the most promising region based on the preliminary hemizygous characterization — in a larger population of male and female 7-month-old homozygous mice and respective WT controls. Importantly, we observed a significant reduction at specific timepoints for all four genes analysed (*Bmal1*, *Clock*, *Per2*, *Cry1*) in homozygous mice compared to WT controls (Fig. 5A-D). Among these, *Bmal1* exhibited the most pronounced dysregulation, with decreases observed at ZT 11 h, ZT 19 h, and in average expression levels across the three timepoints (Fig. 5A). *Clock* and *Cry1* expressions were reduced at ZT 11 h, just before the onset of the night period (Fig. 5C-D), while *Per2* showed a reduction at ZT 3 h, during the light phase (Fig. 5B).

**Figure 5.**
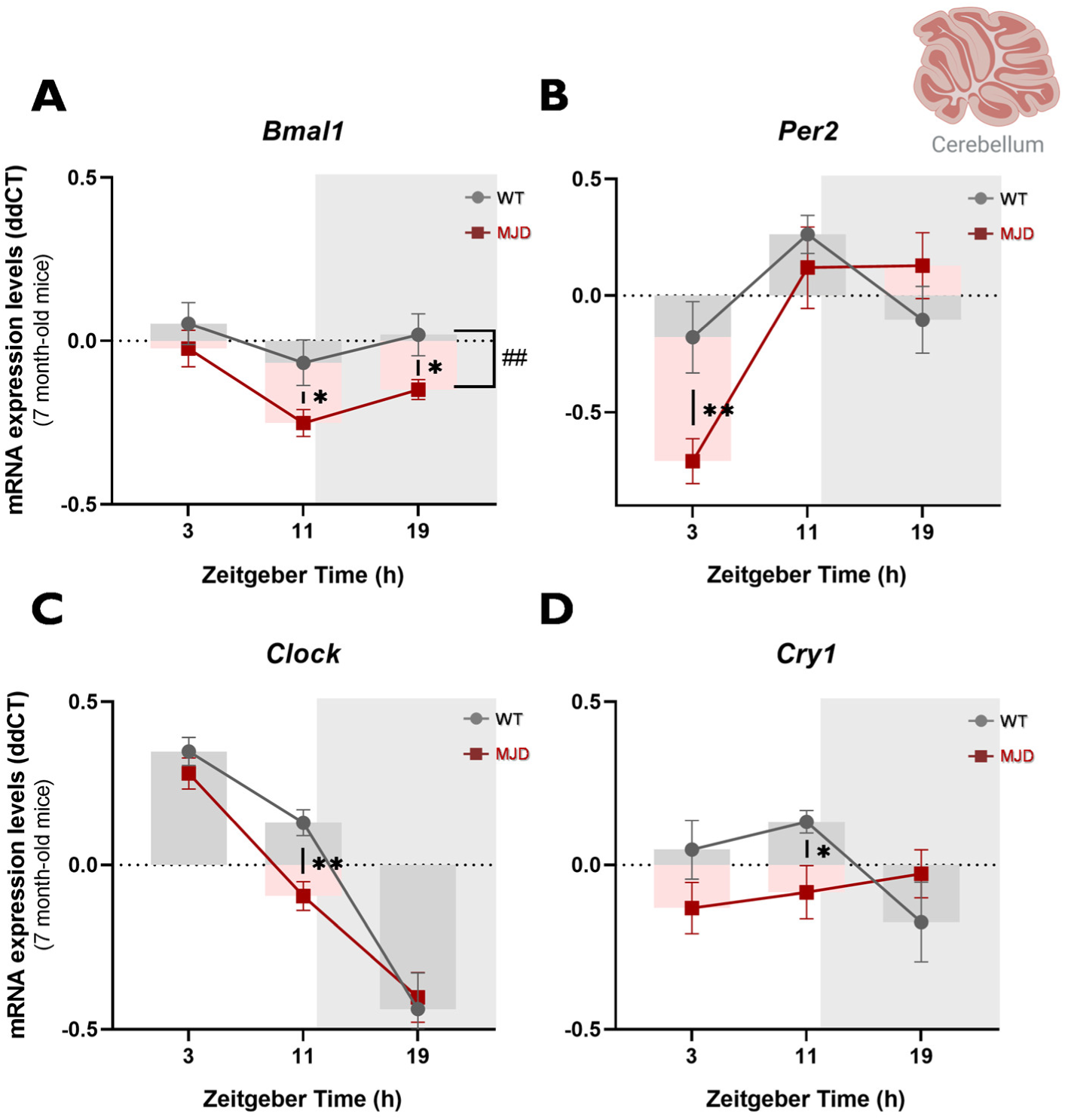
YAC-MJD homozygous mice show a decline in the expression of core clock genes in the cerebellum. Expression levels of 4 core clock genes (A) *Bmal1*, (B) *Per2*, (C) *Clock*, and (D) *Cry1* at 3 timepoints along the day (every 8 hours at zeitgeber time, ZT: 3 h, 11 h, and 19 h) analysed in the cerebellum of 7-month-old MJD YAC84.2/84.2 (homozygous) and wild-type (WT) control mice (*n* = 4-5 mice per timepoint/sex/genotype). Dramatic decreases observed in specific timepoints of all the analysed genes in the homozygous mice when compared to WT controls. (A) *Bmal1* exhibited the most significant dysregulation, with decreases observed at ZT 11 h, ZT 19 h, and in average expression levels for the 3 timepoints. Expression levels of target genes are shown relative to the reference gene *Hprt*. Data are presented as mean ΔΔCT values ± SEM. Grey rectangles indicate the dark phases. Unpaired t tests were performed to assess differences in clock gene expression between homozygous and WT control mice at each timepoint of the day (*, **) and in average expression levels (##). Statistical significance was set as: \**P* < 0.05, \*\**P* < 0.01, ##*P* < 0.01.

Altogether, these findings suggest that clock gene expression is perturbed in the hypothalamus, and, more notably, in the cerebellum of YAC-MJD transgenic mice.

### WT ATXN3 activates the promoter of *Bmal1* and *Per2*, while mutATXN3 loses the capacity to drive *Per2* upon polyglutamine expansion

Given the significant disruption of the clock system observed in the YAC-MJD transgenic mouse model, we investigated whether WT ATXN3 and mutATXN3 interfere with the promoters of the main core clock genes *Bmal1* and *Per2*.

To explore this, we co-transfected N2a cells with *Bmal1*-Luc or *Per2*-Luc reporter plasmids and either mutATXN3 (72 CAG repeats) or WT ATXN3 (27 CAG repeats) constructs. (Fig. 6A). Bioluminescence readings revealed that both WT ATXN3 and mutATXN3 significantly and robustly activate the promoter of *Bmal1* (more than 6-fold increase in bioluminescence compared with cells transfected only with the luciferase reporter; Fig. 6B). On the other hand, only WT ATXN3 significantly increased the transcription of *Per2* (3-fold increment compared with cells transfected only with *Per2*-Luc; Fig. 6C).

**Figure 6.**
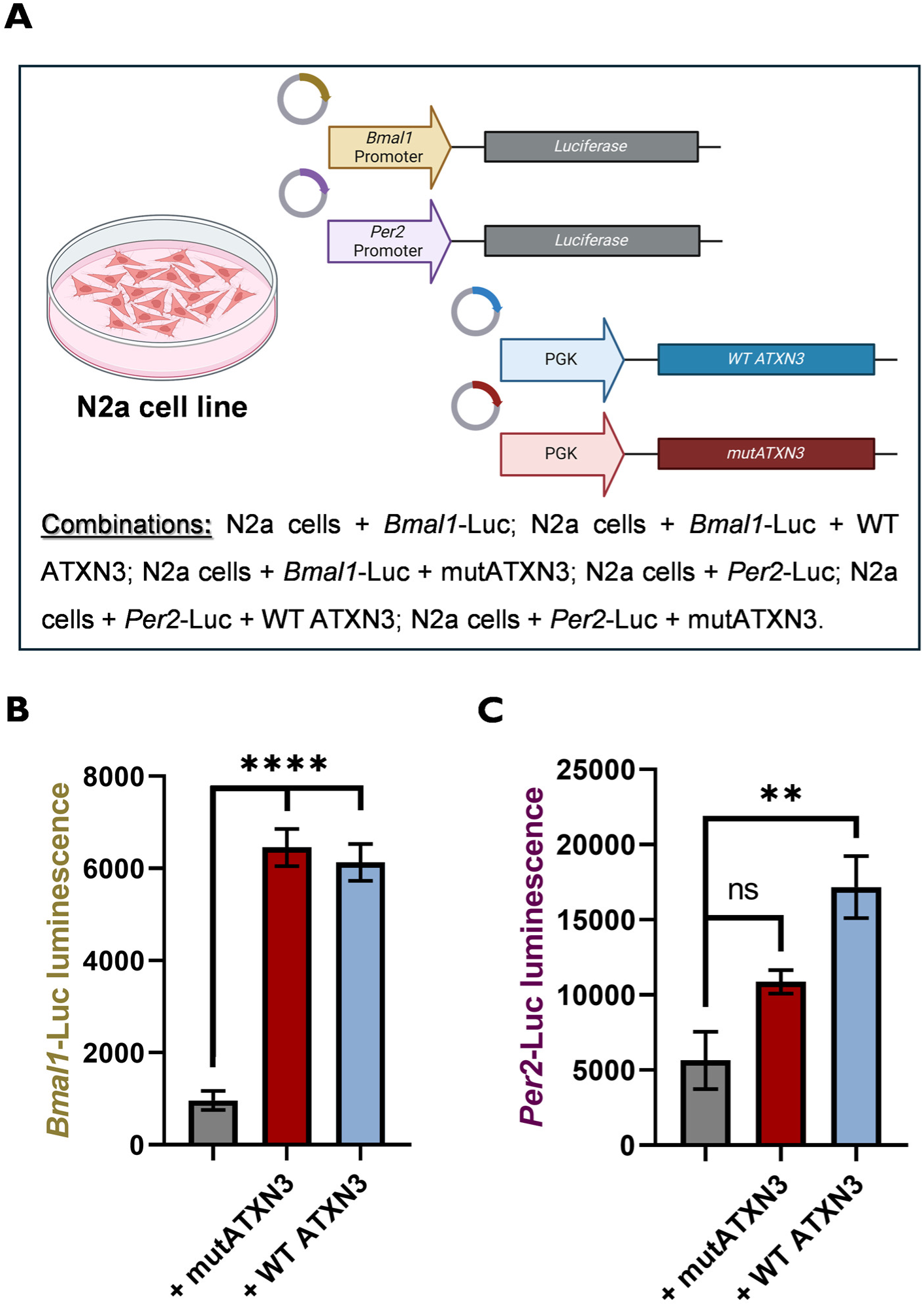
WT ATXN3 activates the promoter of *Bmal1* and *Per2*, while mutATXN3 loses the capacity to drive *Per2* upon polyglutamine expansion. (A) Schematic representation of the experimental conditions. N2a cells were co-transfected with *Bmal1*-Luc or *Per2*-Luc reporter plasmids and human constructs of either mutATXN3 (72 CAG repeats) or WT ATXN3 (27 CAG repeats). Cells were harvested for luciferase assays 24 h after synchronization with 50 % horse serum shock. Analysis of interaction with (B) *Bmal1* and (C) *Per2* promoters by measurement of luciferase bioluminescence in N2a cells. Both WT and mutant forms of ATXN3 drive the promoter of *Bmal1*; however, only the WT ATXN3 form can drive the transcription of *Per2*, a capacity that is lost upon polyglutamine expansion in mutATXN3. (B, C) Values are expressed as mean ± SEM of *n* = 4. Values between groups are compared with one-way ANOVA followed by Dunnett’s multiple comparisons test. Statistical significance was set as: \*\**P* < 0.01, \*\*\*\**P* < 0.0001. ns: not significant.

These observations suggest that the human ATXN3 interacts, directly or indirectly, with the clock machinery, but its capacity to activate the *Per2* promoter is impaired by the polyglutamine expansion in mutATXN3.

## Discussion

Unlike other neurodegenerative disorders, the relationship between the circadian clock and MJD has remained largely unexplored. In this study, we provide the first clinical evidence that rest-activity rhythms are disrupted in MJD patients. We further investigated the specific impairments of circadian rhythms and, importantly, the neurobiological mechanisms underlying clock dysregulation in MJD using the YAC-MJD transgenic mouse model. To assess clock robustness, we evaluated activity patterns under different light schemes, jet-lag re-entrainment capacity, and CBT rhythms. Notably, our findings indicate that the observed circadian dysfunctions may arise from the effects of mutATXN3 on essential neuropeptides within the central clock system and consequent disruptions in core clock gene expression in key brain regions. Additionally, we explored the impact of mutATXN3 on the regulation of core clock genes, such as *Bmal1* and *Per2*, *in vitro*.

Actigraphy enables objective and longitudinal measurement of motor activity in humans within natural living environments using a wristband device.^69^ While it has been widely applied to evaluate sleep-wake alterations, there is growing recognition of its potential in assessing circadian rhythm integrity through validated and automated measures.^69–71^ Accordingly, it has also been used to investigate circadian rhythm disruptions among patients with neurodegenerative diseases; however, its application in MJD has thus far been limited to studying sleep patterns in comparison with polysomnography.^13,23^

In this pilot actigraphy study involving seven MJD patients, we observed a disruption in rest-activity rhythms that showed to be significantly correlated with the clinical progression of MJD. In the course of the disease, patients show decreased rhythm stability (ISm), higher fragmentation of the rest-activity patterns (IVm), and higher activity during the night period (L5). These observations are in line with previous studies showing similar alterations in other neurodegenerative diseases, including Alzheimer’s, Parkinson’s, and Huntington’s diseases.^23,72–74^ Importantly, the increase in nocturnal activity observed in MJD patients appears to be associated with involuntary movements during sleep, as the number of awakenings did not change along the disease course, similar to what has been reported in Huntington’s disease patients.^75^ CFI is an integrative parameter that reflects the robustness of behavioural circadian rhythms, and we found that it has a very strong negative correlation with the SARA score, a well-established clinical scale to assess cerebellar ataxia in MJD.^33,34,40^ This observation reinforces the evidence for a circadian rhythm decline as the disease progresses. Moreover, this finding was also corroborated by other clinical scales (ADL, INAS, and CCFS), highlighting the potential of actigraphy and CFI, as an accurate, personalized, and cost-effective non-invasive digital biomarker of disease progression for MJD. Additionally, we found that amplitude and acrophase tend to decrease with MJD progression. Importantly, lower circadian amplitude and phase advance have been associated with a greater cognitive decline in several neurodegenerative conditions.^23,76^ Furthermore, we observed that both sleep efficiency and total sleep time decreased with the course of MJD.

This pilot study not only provides crucial new insights into the pathophysiology of MJD but also demonstrates, for the first time, the potential of CFI as a biomarker for the progression of a neurodegenerative disease. Given these results, CFI should be further explored in a larger cohort of patients with MJD and in the context of other neurodegenerative pathologies. In the case of MJD, its low prevalence often poses a challenge for larger clinical studies. Despite this, we believe these results warrant further efforts to recruit more patients, allowing more comprehensive analysis and deeper investigation.

To investigate the neurobiological mechanisms underlying the progressive circadian dysfunction in MJD patients, we investigated the circadian rhythms of a YAC-MJD transgenic mouse model. YAC-MJD transgenic mice recapitulate better the disease in MJD patients compared to other mouse models, as they carry the full human mut*ATXN3* gene, including its regulatory, intronic, and promoter regions.^32^

Our initial question was whether circadian activity would also be affected in this mouse model of MJD. A recent study showed that sleep architecture is altered, and motor activity differs in the homozygous mice compared to controls.^26^ However, continuous activity recording or abrupt changes in light schemes were not used to assess a circadian phenotype. Consistently, we observed a motor impairment over time in homozygous mice by conducting, for the first time, wheel-running experiments to continuously record their activity. We found a decline in activity levels under different light schemes (LD and DD) and during the inactive period. Importantly, this difference in total activity was restricted to the early-night period. Homozygous mice also showed increased behavioural fragmentation, similar to what has been reported in the Huntington’s disease transgenic mouse model BACHD (expressing the entire human *HTT* gene).^57^

Current evidence suggests that aligning behavioural rhythms to acute phase shifts in the environmental LD cycle is a key function of the circadian system, with jet lag protocols being used to evaluate this capacity.^31,56,57^ One of the major findings of our study is that the homozygous mice take more than double the time to fully adapt to a 4 h phase advance, compared to WT controls. For reference, this difference exceeds that observed in mouse models of Huntington’s disease and is equivalent to that seen in brain-specific SIRT1 knockout mice.^31,56,57^ This pronounced dysfunction was confirmed in older hemizygous mice, which took longer to re-entrain to the new LD cycle, albeit with a reduced percentage of increase, compared to their age-matched controls. Notably, 7-month-old homozygous mice needed even more time to re-synchronize than 13-month-old WT mice. The number of days to adapt in both cohorts of WT mice is in accordance with what has been described in previous studies and further confirms the decline in central circadian function with ageing.^31^ While further experiments are needed (e.g. in larger cohorts, in female mice, and evaluating LD phase delays), the observed circadian dysfunction is in line with the actigraphy data and confirms the decline in circadian robustness in MJD.

In addition to behavioural rest-activity rhythms, biological biomarkers such as CBT are also directly controlled by the master clock and well-accepted markers to investigate potential circadian disruptions. According to Colwell^22^, CBT should be evaluated more frequently as a measure of SCN-driven output in mouse models of neurodegenerative diseases.^22,23^ We found that homozygous mice have a higher CBT compared to WT mice, more pronounced at the onset of their active phase. These findings are aligned with preliminary measurements in the MJD patients included in this study, which show increased MESOR values of skin temperature relative to healthy individuals (data now shown). Increased CBT has been reported in other neurodegenerative models.^57,77^ In Huntington’s disease, CBT is approximately 0.3 °C higher in the BACHD mouse model, particularly during the inactive phase, and in patients even decades before disease onset.^57,78,79^ This increase in CBT and sympathetic tone is hypothesized to be associated with a central autonomic network dysfunction, which has also been extensively reported in MJD patients.^80^ Furthermore, when analysing the circadian parameters of CBT, we observed that homozygous mice exhibit an increase in MESOR and amplitude, and a phase-advance of over 1 h in the CBT peak. These findings are consistent with the potential phase advance observed in the rest-activity cycles of patients with MJD as the disease progresses. Similar changes in amplitude and acrophase have been reported in the 3xTgAD Alzheimer’s disease mouse model in several studies.^77,81^ Interestingly, patients with mild cognitive impairment present a phase-advance in both activity and skin temperature rhythm.^82^ The similarities in CBT dysregulation between models of other neurodegenerative diseases and our findings in MJD suggest a potential mechanistic overlap. However, further investigations into CBT circadian rhythms in MJD patients and other markers associated with thermoregulation in the YAC-MJD mouse model, such as brown-adipose tissue thermogenesis, are needed to better understand the observed alterations.

It has been hypothesized that the observed alterations in CBT circadian parameters are, at least partially, caused by a reduction in the neuropeptides VIP and AVP in the hypothalamic SCN and PVN.^81^ Accordingly, we observed a reduction of VIP and AVP levels in the SCN of homozygous mice. The 3xTgAD mouse model also presents decreased levels of VIP and AVP in the SCN, even before the pathology onset, which is suggested as the mechanistic reason behind the increased amplitude and phase-advance observed in the CBT of these mice.^81,83^ The decreased levels of these neuropeptides in the SCN have also been well documented in Alzheimer’s disease *postmortem* samples.^84,85^ Further characterization in Alzheimer’s disease patients measured activity rhythms and examined their SCN *postmortem*, revealing that the loss of AVP was associated with increased activity fragmentation, while the loss of VIP correlated with a dampened amplitude of motor activity.^58,59^ The decrease in the levels of VIP and AVP was also observed in mouse models (BACHD) and patients with Huntington’s disease.^60,61^ It has been suggested that circadian rhythm disturbances arise from pathology of the SCN^61^ and, in fact, SCN projections to the PVN are essential to regulate circadian rhythms, including core body temperature, and to synchronize the peripheral clocks.^63,65,67^ We found a dramatic 78 % reduction in the number of AVP positive cells in the PVN of homozygous mice compared to controls, even greater than the 65 % reduction observed in the SCN. These findings suggest that neuronal alterations in the PVN may also contribute to the circadian disturbances observed in MJD. Furthermore, electrophysiological studies of the SCN and PVN in the YAC-MJD mouse model, and evaluation of VIP and AVP in the master clock of MJD patients are two important future steps to further understand these dysfunctions in MJD.

Given the importance of central clock synchronization for the robustness of the circadian clock system, we also assessed the expression levels of the core clock genes *Bmal1*, *Clock*, *Per2*, and *Cry1* in the hypothalamus and cerebellum of YAC-MJD mice every 8 h.^64^ Core clock genes are the building blocks of the clock system at the molecular level. The proteins they encode act as transcription factors which further regulate the expression of multiple other genes (clock-controlled genes), involved in various physiological processes (e.g., metabolism, hormone regulation, immune response, and sleep) including neuronal functions.^15,17,54^

Importantly, we found a significant reduction in the expression levels of *Bmal1*, *Clock*, *Per2*, and *Cry1* at specific timepoints throughout the day in the cerebellum of homozygous mice, a brain region heavily affected in MJD and characterized by pronounced gene expression rhythms. Reduced expression of core clock genes has been reported in several studies on other neurodegenerative disorders, including Parkinson’s, Alzheimer’s, and Huntington’s diseases.^86^ Surprisingly, no changes in the molecular clock were found in the BACHD mouse model.^57,87^ Others reported impairment of *BMAL1* and *CLOCK* expression in human induced pluripotent stem cells derived from patients with SCA17; however, no further details were provided (e.g., increase/decrease or number of timepoints assessed).^88^ One of the major findings of our study is the decline of *Bmal1* expression at ZT 11 h, ZT 19 h, and in average expression levels in the cerebellum of homozygous mice, suggesting a strong impact of the disease on the clock molecular machinery. Conditional *Bmal1* knock-out in cerebellar Purkinje cells was shown to induce behavioural and cellular changes that are even more pronounced compared to a *Bmal1* knock-out mice, reinforcing the significance of our findings.^28^ Experiments in larger cohorts, with more timepoints, in isolated regions of the hypothalamus (e.g. in SCN), and other organs should be performed in order to better characterize clock gene expression in MJD. It would also be relevant to evaluate clock gene expression in peripheral blood mononuclear cells of MJD patients and correlate with actigraphy results.

The multilevel dysfunctions observed in the YAC-MJD transgenic mouse model suggested that the expression of mutATXN3 could interfere, directly or not, with the normal regulation of clock gene expression. Accordingly, we found that WT ATXN3 activates the promoter of *Bmal1* and *Per2*, whereas mutATXN3 loses the capacity to drive *Per2* upon polyglutamine expansion. Interestingly, the higher magnitude of increase in *Bmal1* bioluminescence, compared to *Per2*, after transfection with WT ATXN3 was similar to what has been observed for SIRT1, a known strong regulator of the clock.^31^ Previous reports have shown that *Atxn3* expression exhibits an oscillating profile; however, the impact on the clock molecular machinery was not investigated before.^27^ More remarkably, *Bmal1* knock-out mice revealed a strong decrease in the cerebellar expression of *Atxn3,* suggesting a common pathway.^28^ Our findings reinforce and expand upon these observations, by demonstrating the robust effect of WT ATXN3 in the promoter of *Bmal1* and *Per2*. Furthermore, we show that the polyglutamine expansion of *ATXN3* in MJD disrupts its normal direct or indirect interaction with clock genes, particularly *Per2*, which is likely to reflect on downstream pathways. Accordingly, previous studies have shown that *Per2* deficiencies in mice are linked to alterations in temperature regulation^89^, lipid metabolism^90^, and behavioural re-entrainment during jet lag^91^.

Altogether, these findings offer a potential mechanistic explanation for the circadian alterations observed in MJD. To the best of our knowledge, this is the first study showing that clock regulation is markedly affected in MJD patients and in both *in vivo* and *in vitro* models of the disease. Understanding clock dysfunction in MJD is crucial not only to better understand MJD pathophysiology but also to provide preliminary foundations for the identification of biomarkers to monitor disease progression (e.g. actigraphy parameters, clock gene expression, CBT analysis) and to test circadian-based interventions to tackle this devastating disease.^17,87^ Lastly, our study reinforces that circadian rhythms must be taken into account in neurodegenerative diseases’ research and clinical management, including MJD and other SCAs.

## Data availability

The data that support the findings of this study are available from the corresponding author, upon request.

## Acknowledgements

We would like to thank the CNC-UC animal facility, in particular Paula Mota and Daniela Talhada for the help with the implantation of the telemetric data loggers. We thank Luisa Cortes, Margarida Caldeira, Tatiana Catarino, and the CNC MICC team for assistance with microscopy imaging and image analysis. *Bmal1/Per2*-LUC plasmids were kindly provided by Angela Relógio. *Hprt* primer was kindly provided by Rui Jorge Nobre. We thank Rosário Faro for the support with plasmid preparation. We thank Jisette Gonzalez for the help with plate coating and the Bright-Glo^®^ protocol. We thank all members of the L.P.d.A. and C.C. labs for all the support, discussion, and comments. We thank the previous members of C.C. lab, Sara Silva and Célia Aveleira, for their support and technical expertise in the initial steps of this study. Schematic figures were created using BioRender.com.

## References

1. Schöls L, Bauer P, Schmidt T, Schulte T, Riess O. Autosomal dominant cerebellar ataxias: clinical features, genetics, and pathogenesis. Lancet Neurol. 2004;3(5):291–304. doi:10.1016/S1474-4422(04)00737-9

2. Takiyama Y, Nishizawa M, Tanaka H, et al. The gene for Machado-Joseph disease maps to human chromosome 14q. Nat Genet. 1993;4(3):300–304. doi:10.1038/ng0793-300

3. Paulson HL, Das SS, Crino PB, et al. Machado-Joseph disease gene product is a cytoplasmic protein widely expressed in brain. Ann Neurol. 1997;41(4):453–462. doi:10.1002/ana.410410408

4. Kawaguchi Y, Okamoto T, Taniwaki M, et al. CAG expansions in a novel gene for Machado-Joseph disease at chromosome 14q32.1. Nat Genet. 1994;8(3):221–228. doi:10.1038/ng1194-221

5. Klockgether T, Skalej M, Wedekind D, et al. Autosomal dominant cerebellar ataxia type I. MRI-based volumetry of posterior fossa structures and basal ganglia in spinocerebellar ataxia types 1, 2 and 3. Brain. 1998;121 (Pt 9):1687–1693. doi:10.1093/brain/121.9.1687

6. Ikeda H, Yamaguchi M, Sugai S, Aze Y, Narumiya S, Kakizuka A. Expanded polyglutamine in the Machado-Joseph disease protein induces cell death in vitro and in vivo. Nat Genet. 1996;13(2):196–202. doi:10.1038/ng0696-196

7. Maruyama H, Nakamura S, Matsuyama Z, et al. Molecular features of the CAG repeats and clinical manifestation of Machado-Joseph disease. Hum Mol Genet. 1995;4(5):807–812. doi:10.1093/hmg/4.5.807

8. Schöls L, Bauer P, Schmidt T, Schulte T, Riess O. Autosomal dominant cerebellar ataxias: clinical features, genetics, and pathogenesis. Lancet Neurol. 2004;3(5):291–304. doi:10.1016/S1474-4422(04)00737-9

9. Coutinho P, Andrade C. Autosomal dominant system degeneration in Portuguese families of the Azores Islands. A new genetic disorder involving cerebellar, pyramidal, extrapyramidal and spinal cord motor functions. Neurology. 1978;28(7):703–709. doi:10.1212/wnl.28.7.703

10. Matos CA, de Almeida LP, Nóbrega C. Machado-Joseph disease/spinocerebellar ataxia type 3: lessons from disease pathogenesis and clues into therapy. J Neurochem. 2019;148(1):8–28. doi:10.1111/jnc.14541

11. Cunha-Santos J, Duarte-Neves J, Carmona V, Guarente L, Pereira de Almeida L, Cavadas C. Caloric restriction blocks neuropathology and motor deficits in Machado-Joseph disease mouse models through SIRT1 pathway. Nat Commun. 2016;7:11445. doi:10.1038/ncomms11445

12. Pedroso JL, Braga-Neto P, Felício AC, et al. Sleep disorders in machado-joseph disease: frequency, discriminative thresholds, predictive values, and correlation with ataxia-related motor and non-motor features. Cerebellum. 2011;10(2):291–295. doi:10.1007/s12311-011-0252-7

13. LaGrappe D, Massey L, Kruavit A, et al. Sleep disorders among Aboriginal Australians with Machado-Joseph Disease: Quantitative results from a multiple methods study to assess the experience of people living with the disease and their caregivers. Neurobiol Sleep Circadian Rhythms. 2022;12:100075. doi:10.1016/j.nbscr.2022.100075

14. Folha Santos FA, de Carvalho LBC, Prado LF do, do Prado GF, Barsottini OG, Pedroso JL. Sleep apnea in Machado-Joseph disease: a clinical and polysomnographic evaluation. Sleep Med. 2018;48:23–26. doi:10.1016/j.sleep.2018.04.002

15. Ribeiro RFN, Cavadas C, Silva MMC. Small-molecule modulators of the circadian clock: Pharmacological potentials in circadian-related diseases. Drug Discov Today. 2021;26(7):1620–1641. doi:10.1016/j.drudis.2021.03.015

16. Rijo-Ferreira F, Takahashi JS. Genomics of circadian rhythms in health and disease. Genome Med. 2019;11(1):82. doi:10.1186/s13073-019-0704-0

17. Ribeiro RFN, Pereira D, de Almeida LP, Silva MMC, Cavadas C. SIRT1 activation and its circadian clock control: a promising approach against (frailty in) neurodegenerative disorders. Aging Clin Exp Res. 2022;34(12):2963–2976. doi:10.1007/s40520-022-02257-y

18. Hood S, Amir S. The aging clock: circadian rhythms and later life. J Clin Invest. 2017;127(2):437–446. doi:10.1172/JCI90328

19. Bass J, Lazar MA. Circadian time signatures of fitness and disease. Science. 2016;354(6315):994–999. doi:10.1126/science.aah4965

20. Liu AC, Lewis WG, Kay SA. Mammalian circadian signaling networks and therapeutic targets. Nat Chem Biol. 2007;3(10):630–639. doi:10.1038/nchembio.2007.37

21. Hood S, Amir S. Neurodegeneration and the Circadian Clock. Front Aging Neurosci. 2017;9:170. doi:10.3389/fnagi.2017.00170

22. Colwell CS. Defining circadian disruption in neurodegenerative disorders. J Clin Invest. 2021;131(19). doi:10.1172/JCI148288

23. Leng Y, Musiek ES, Hu K, Cappuccio FP, Yaffe K. Association between circadian rhythms and neurodegenerative diseases. Lancet Neurol. 2019;18(3):307–318. doi:10.1016/S1474-4422(18)30461-7

24. Nassan M, Videnovic A. Circadian rhythms in neurodegenerative disorders. Nat Rev Neurol. 2022;18(1):7–24. doi:10.1038/s41582-021-00577-7

25. Shen Y, Lv QK, Xie WY, et al. Circadian disruption and sleep disorders in neurodegeneration. Transl Neurodegener. 2023;12(1):8. doi:10.1186/s40035-023-00340-6

26. Tsimpanouli ME, Ghimire A, Barget AJ, et al. Sleep Alterations in a Mouse Model of Spinocerebellar Ataxia Type 3. Cells. 2022;11(19). doi:10.3390/cells11193132

27. Zhang R, Lahens NF, Ballance HI, Hughes ME, Hogenesch JB. A circadian gene expression atlas in mammals: implications for biology and medicine. Proc Natl Acad Sci U S A. 2014;111(45):16219–16224. doi:10.1073/pnas.1408886111

28. Liu D, Nanclares C, Simbriger K, et al. Autistic-like behavior and cerebellar dysfunction in Bmal1 mutant mice ameliorated by mTORC1 inhibition. Mol Psychiatry. 2023;28(9):3727–3738. doi:10.1038/s41380-022-01499-6

29. Watanave M, Hoshino C, Konno A, et al. Pharmacological enhancement of retinoid-related orphan receptor α function mitigates spinocerebellar ataxia type 3 pathology. Neurobiol Dis. 2019;121:263–273. doi:10.1016/j.nbd.2018.10.014

30. Yasui H, Matsuzaki Y, Konno A, Hirai H. Global Knockdown of Retinoid-related Orphan Receptor α in Mature Purkinje Cells Reveals Aberrant Cerebellar Phenotypes of Spinocerebellar Ataxia. Neuroscience. 2021;462:328–336. doi:10.1016/j.neuroscience.2020.04.004

31. Chang HC, Guarente L. SIRT1 mediates central circadian control in the SCN by a mechanism that decays with aging. Cell. 2013;153(7):1448–1460. doi:10.1016/j.cell.2013.05.027

32. Cemal CK, Carroll CJ, Lawrence L, et al. YAC transgenic mice carrying pathological alleles of the MJD1 locus exhibit a mild and slowly progressive cerebellar deficit. Hum Mol Genet. 2002;11(9):1075–1094. doi:10.1093/hmg/11.9.1075

33. Schmitz-Hübsch T, du Montcel ST, Baliko L, et al. Scale for the assessment and rating of ataxia: development of a new clinical scale. Neurology. 2006;66(11):1717–1720. doi:10.1212/01.wnl.0000219042.60538.92

34. Lima M, Raposo M, Ferreira A, et al. The Homogeneous Azorean Machado-Joseph Disease Cohort: Characterization and Contributions to Advances in Research. Biomedicines. 2023;11(2). doi:10.3390/biomedicines11020247

35. Jacobi H, Rakowicz M, Rola R, et al. Inventory of Non-Ataxia Signs (INAS): validation of a new clinical assessment instrument. Cerebellum. 2013;12(3):418–428. doi:10.1007/s12311-012-0421-3

36. du Montcel ST, Charles P, Ribai P, et al. Composite cerebellar functional severity score: validation of a quantitative score of cerebellar impairment. Brain. 2008;131(Pt 5):1352–1361. doi:10.1093/brain/awn059

37. Edemekong PF, Bomgaars DL, Sukumaran S, Schoo C. Activities of Daily Living.; 2024.

38. Buysse DJ, Reynolds CF, Monk TH, Berman SR, Kupfer DJ. The Pittsburgh Sleep Quality Index: a new instrument for psychiatric practice and research. Psychiatry Res. 1989;28(2):193–213. doi:10.1016/0165-1781(89)90047-4

39. Gubin D, Danilenko K, Stefani O, et al. Blue Light and Temperature Actigraphy Measures Predicting Metabolic Health Are Linked to Melatonin Receptor Polymorphism. Biology (Basel). 2023;13(1). doi:10.3390/biology13010022

40. Ortiz-Tudela E, Martinez-Nicolas A, Campos M, Rol MÁ, Madrid JA. A new integrated variable based on thermometry, actimetry and body position (TAP) to evaluate circadian system status in humans. PLoS Comput Biol. 2010;6(11):e1000996. doi:10.1371/journal.pcbi.1000996

41. Witting W, Kwa IH, Eikelenboom P, Mirmiran M, Swaab DF. Alterations in the circadian rest-activity rhythm in aging and Alzheimer’s disease. Biol Psychiatry. 1990;27(6):563–572. doi:10.1016/0006-3223(90)90523-5

42. Gonçalves BSB, Cavalcanti PRA, Tavares GR, Campos TF, Araujo JF. Nonparametric methods in actigraphy: An update. Sleep Sci. 2014;7(3):158–164. doi:10.1016/j.slsci.2014.09.013

43. Cornelissen G. Cosinor-based rhythmometry. Theor Biol Med Model. 2014;11:16. doi:10.1186/1742-4682-11-16

44. Pritchett D, Jagannath A, Brown LA, et al. Deletion of Metabotropic Glutamate Receptors 2 and 3 (mGlu2 & mGlu3) in Mice Disrupts Sleep and Wheel-Running Activity, and Increases the Sensitivity of the Circadian System to Light. PLoS One. 2015;10(5):e0125523. doi:10.1371/journal.pone.0125523

45. Rufiange M, Leung VSY, Simpson K, Pang DSJ. Pre-warming before general anesthesia with isoflurane delays the onset of hypothermia in rats. PLoS One. 2020;15(3):e0219722. doi:10.1371/journal.pone.0219722

46. Mai TC, Delanaud S, Bach V, Braun A, Pelletier A, de Seze R. Effect of non-thermal radiofrequency on body temperature in mice. Sci Rep. 2020;10(1):5724. doi:10.1038/s41598-020-62789-z

47. Parsons R, Parsons R, Garner N, Oster H, Rawashdeh O. CircaCompare: a method to estimate and statistically support differences in mesor, amplitude and phase, between circadian rhythms. Bioinformatics. 2020;36(4):1208–1212. doi:10.1093/bioinformatics/btz730

48. Wu Z, Martinez ME, Hernandez A. Mice lacking DIO3 exhibit sex-specific alterations in circadian patterns of corticosterone and gene expression in metabolic tissues. BMC Mol Cell Biol. 2024;25(1):11. doi:10.1186/s12860-024-00508-6

49. Huang R, Chen J, Zhou M, et al. Multi-omics profiling reveals rhythmic liver function shaped by meal timing. Nat Commun. 2023;14(1):6086. doi:10.1038/s41467-023-41759-9

50. McNeill JK, Walton JC, Ryu V, Albers HE. The Excitatory Effects of GABA within the Suprachiasmatic Nucleus: Regulation of Na-K-2Cl Cotransporters (NKCCs) by Environmental Lighting Conditions. J Biol Rhythms. 2020;35(3):275–286. doi:10.1177/0748730420924271

51. Alves S, Régulier E, Nascimento-Ferreira I, et al. Striatal and nigral pathology in a lentiviral rat model of Machado-Joseph disease. Hum Mol Genet. 2008;17(14):2071–2083. doi:10.1093/hmg/ddn106

52. Carmo-Silva S, Ferreira-Marques M, Nóbrega C, et al. Ataxin-2 in the hypothalamus at the crossroads between metabolism and clock genes. J Mol Endocrinol. 2023;70(1). doi:10.1530/JME-21-0272

53. Livak KJ, Schmittgen TD. Analysis of relative gene expression data using real-time quantitative PCR and the 2(-Delta Delta C(T)) Method. Methods. 2001;25(4):402–408. doi:10.1006/meth.2001.1262

54. Gaspar LS, Hesse J, Yalçin M, et al. Long-term continuous positive airway pressure treatment ameliorates biological clock disruptions in obstructive sleep apnea. EBioMedicine. 2021;65:103248. doi:10.1016/j.ebiom.2021.103248

55. Gao C, Haghayegh S, Wagner M, et al. Approaches for Assessing Circadian Rest-Activity Patterns Using Actigraphy in Cohort and Population-Based Studies. Curr Sleep Med Rep. 2023;9(4):247–256. doi:10.1007/s40675-023-00267-4

56. Wood NI, McAllister CJ, Cuesta M, Aungier J, Fraenkel E, Morton AJ. Adaptation to experimental jet-lag in R6/2 mice despite circadian dysrhythmia. PLoS One. 2013;8(2):e55036. doi:10.1371/journal.pone.0055036

57. Kudo T, Schroeder A, Loh DH, et al. Dysfunctions in circadian behavior and physiology in mouse models of Huntington’s disease. Exp Neurol. 2011;228(1):80–90. doi:10.1016/j.expneurol.2010.12.011

58. Harper DG, Stopa EG, Kuo-Leblanc V, et al. Dorsomedial SCN neuronal subpopulations subserve different functions in human dementia. Brain. 2008;131(Pt 6):1609–1617. doi:10.1093/brain/awn049

59. Wang JL, Lim AS, Chiang WY, et al. Suprachiasmatic neuron numbers and rest-activity circadian rhythms in older humans. Ann Neurol. 2015;78(2):317–322. doi:10.1002/ana.24432

60. Kuljis DA, Gad L, Loh DH, et al. Sex Differences in Circadian Dysfunction in the BACHD Mouse Model of Huntington’s Disease. PLoS One. 2016;11(2):e0147583. doi:10.1371/journal.pone.0147583

61. van Wamelen DJ, Aziz NA, Anink JJ, et al. Suprachiasmatic nucleus neuropeptide expression in patients with Huntington’s Disease. Sleep. 2013;36(1):117–125. doi:10.5665/sleep.2314

62. Perreau-Lenz S, Kalsbeek A, Pévet P, Buijs RM. Glutamatergic clock output stimulates melatonin synthesis at night. Eur J Neurosci. 2004;19(2):318–324. doi:10.1111/j.0953-816x.2003.03132.x

63. Lu J, Zhang YH, Chou TC, et al. Contrasting effects of ibotenate lesions of the paraventricular nucleus and subparaventricular zone on sleep-wake cycle and temperature regulation. J Neurosci. 2001;21(13):4864–4874. doi:10.1523/JNEUROSCI.21-13-04864.2001

64. Starnes AN, Jones JR. Inputs and Outputs of the Mammalian Circadian Clock. Biology (Basel). 2023;12(4). doi:10.3390/biology12040508

65. Paul S, Hanna L, Harding C, et al. Output from VIP cells of the mammalian central clock regulates daily physiological rhythms. Nat Commun. 2020;11(1):1453. doi:10.1038/s41467-020-15277-x

66. Abrahamson EE, Moore RY. Suprachiasmatic nucleus in the mouse: retinal innervation, intrinsic organization and efferent projections. Brain Res. 2001;916(1-2):172–191. doi:10.1016/s0006-8993(01)02890-6

67. Jones JR, Chaturvedi S, Granados-Fuentes D, Herzog ED. Circadian neurons in the paraventricular nucleus entrain and sustain daily rhythms in glucocorticoids. Nat Commun. 2021;12(1):5763. doi:10.1038/s41467-021-25959-9

68. Mendoza J, Pévet P, Felder-Schmittbuhl MP, Bailly Y, Challet E. The cerebellum harbors a circadian oscillator involved in food anticipation. J Neurosci. 2010;30(5):1894–1904. doi:10.1523/JNEUROSCI.5855-09.2010

69. Ali FZ, Parsey R V, Lin S, Schwartz J, DeLorenzo C. Circadian rhythm biomarker from wearable device data is related to concurrent antidepressant treatment response. NPJ Digit Med. 2023;6(1):81. doi:10.1038/s41746-023-00827-6

70. Ancoli-Israel S, Cole R, Alessi C, Chambers M, Moorcroft W, Pollak CP. The role of actigraphy in the study of sleep and circadian rhythms. Sleep. 2003;26(3):342–392. doi:10.1093/sleep/26.3.342

71. Fossion R, Rivera AL, Toledo-Roy JC, Ellis J, Angelova M. Multiscale adaptive analysis of circadian rhythms and intradaily variability: Application to actigraphy time series in acute insomnia subjects. PLoS One. 2017;12(7):e0181762. doi:10.1371/journal.pone.0181762

72. Harper DG, Stopa EG, McKee AC, Satlin A, Fish D, Volicer L. Dementia severity and Lewy bodies affect circadian rhythms in Alzheimer disease. Neurobiol Aging. 2004;25(6):771–781. doi:10.1016/j.neurobiolaging.2003.04.009

73. Morton AJ, Wood NI, Hastings MH, Hurelbrink C, Barker RA, Maywood ES. Disintegration of the sleep-wake cycle and circadian timing in Huntington’s disease. J Neurosci. 2005;25(1):157–163. doi:10.1523/JNEUROSCI.3842-04.2005

74. Musiek ES, Bhimasani M, Zangrilli MA, Morris JC, Holtzman DM, Ju YES. Circadian Rest-Activity Pattern Changes in Aging and Preclinical Alzheimer Disease. JAMA Neurol. 2018;75(5):582–590. doi:10.1001/jamaneurol.2017.4719

75. Hurelbrink CB, Lewis SJG, Barker RA. The use of the Actiwatch-Neurologica system to objectively assess the involuntary movements and sleep-wake activity in patients with mild-moderate Huntington’s disease. J Neurol. 2005;252(6):642–647. doi:10.1007/s00415-005-0709-z

76. Rogers-Soeder TS, Blackwell T, Yaffe K, et al. Rest-Activity Rhythms and Cognitive Decline in Older Men: The Osteoporotic Fractures in Men Sleep Study. J Am Geriatr Soc. 2018;66(11):2136–2143. doi:10.1111/jgs.15555

77. Tournissac M, Vu TM, Vrabic N, et al. Repurposing beta-3 adrenergic receptor agonists for Alzheimer’s disease: beneficial effects in a mouse model. Alzheimers Res Ther. 2021;13(1):103. doi:10.1186/s13195-021-00842-3

78. Schultz JL, Heinzerling AE, Brinker AN, et al. Autonomic changes in Huntington’s disease correlate with altered central autonomic network connectivity. Brain Commun. 2022;4(5). doi:10.1093/braincomms/fcac253

79. Schultz JL, Harshman LA, Kamholz JA, Nopoulos PC. Autonomic dysregulation as an early pathologic feature of Huntington Disease. Auton Neurosci. 2021;231:102775. doi:10.1016/j.autneu.2021.102775

80. Urbini N, Siciliano L, Olivito G, Leggio M. Unveiling the role of cerebellar alterations in the autonomic nervous system: a systematic review of autonomic dysfunction in spinocerebellar ataxias. J Neurol. 2023;270(12):5756–5772. doi:10.1007/s00415-023-11993-8

81. Knight EM, Brown TM, Gümüsgöz S, et al. Age-related changes in core body temperature and activity in triple-transgenic Alzheimer’s disease (3xTgAD) mice. Dis Model Mech. 2013;6(1):160–170. doi:10.1242/dmm.010173

82. Ortiz-Tudela E, Martinez-Nicolas A, Díaz-Mardomingo C, et al. The characterization of biological rhythms in mild cognitive impairment. Biomed Res Int. 2014;2014:524971. doi:10.1155/2014/524971

83. Sterniczuk R, Dyck RH, Laferla FM, Antle MC. Characterization of the 3xTg-AD mouse model of Alzheimer’s disease: part 1. Circadian changes. Brain Res. 2010;1348:139–148. doi:10.1016/j.brainres.2010.05.013

84. Swaab DF, Fliers E, Partiman TS. The suprachiasmatic nucleus of the human brain in relation to sex, age and senile dementia. Brain Res. 1985;342(1):37–44. doi:10.1016/0006-8993(85)91350-2

85. Zhou JN, Hofman MA, Swaab DF. VIP neurons in the human SCN in relation to sex, age, and Alzheimer’s disease. Neurobiol Aging. 1995;16(4):571–576. doi:10.1016/0197-4580(95)00043-e

86. Canever JB, Queiroz LY, Soares ES, de Avelar NCP, Cimarosti HI. Circadian rhythm alterations affecting the pathology of neurodegenerative diseases. J Neurochem. 2024;168(8):1475–1489. doi:10.1111/jnc.15883

87. Whittaker DS, Loh DH, Wang HB, et al. Circadian-based Treatment Strategy Effective in the BACHD Mouse Model of Huntington’s Disease. J Biol Rhythms. 2018;33(5):535–554. doi:10.1177/0748730418790401

88. F. Motolese, A. Casamassa, A. Vescovi, V. Di Lazzaro, J. Rosati, M. Marano. Circadian rhythm alterations in an in vitro cellular model of Spinocerebellar ataxia type 17. MDS Virtual Congress 2020; Abstract Number: 1149. 2020. Accessed August 22, 2024. https://www.mdsabstracts.org/abstract/circadian-rhythm-alterations-in-an-in-vitro-cellular-model-of-spinocerebellar-ataxia-type-17/

89. Chappuis S, Ripperger JA, Schnell A, et al. Role of the circadian clock gene Per2 in adaptation to cold temperature. Mol Metab. 2013;2(3):184–193. doi:10.1016/j.molmet.2013.05.002

90. Grimaldi B, Bellet MM, Katada S, et al. PER2 controls lipid metabolism by direct regulation of PPARγ. Cell Metab. 2010;12(5):509–520. doi:10.1016/j.cmet.2010.10.005

91. Kiessling S, Eichele G, Oster H. Adrenal glucocorticoids have a key role in circadian resynchronization in a mouse model of jet lag. J Clin Invest. 2010;120(7):2600–2609. doi:10.1172/JCI41192

