## Supplementary for "Circadian Rhythms are Disrupted in Patients and Preclinical Models of Machado-Joseph Disease"

## -   Supplementary material   -

**Supplementary Table 1 – Statistic analysis of YAC-MJD homozygous mice core body temperatures in the beginning of the active dark phase (all significant values presented, related to Figure 3).**

| Time (h:mm:ss) | Mean of WT | Mean of MJD | Difference | Adjusted *P* Value |
|---|---|---|---|---|
| 19:51:40 | 35.83 | 36.59 | -0.7621 | 0.010065 |
| 19:56:39 | 35.81 | 36.67 | -0.8543 | 0.001067 |
| 20:01:39 | 35.82 | 36.74 | -0.9191 | 0.000191 |
| 20:06:39 | 35.79 | 36.79 | -0.9974 | 0.000021 |
| 20:11:39 | 35.84 | 36.84 | -0.994 | 0.000023 |
| 20:16:39 | 35.84 | 36.89 | -1.051 | 0.000004 |
| 20:21:38 | 35.86 | 36.92 | -1.06 | 0.000003 |
| 20:26:38 | 35.95 | 37.03 | -1.078 | 0.000002 |
| 20:31:38 | 36.06 | 37.07 | -1.009 | 0.000015 |
| 20:36:38 | 36.11 | 37.19 | -1.076 | 0.000002 |
| 20:41:37 | 36.19 | 37.17 | -0.9797 | 0.000035 |
| 20:46:37 | 36.2 | 37.18 | -0.9821 | 0.000033 |
| 20:51:37 | 36.27 | 37.23 | -0.9651 | 0.000053 |
| 20:56:37 | 36.3 | 37.23 | -0.9324 | 0.000133 |
| 21:01:37 | 36.27 | 37.22 | -0.9531 | 0.000075 |
| 21:06:36 | 36.3 | 37.24 | -0.936 | 0.00012 |
| 21:11:36 | 36.31 | 37.24 | -0.9253 | 0.000161 |
| 21:16:36 | 36.34 | 37.23 | -0.8957 | 0.000359 |
| 21:21:36 | 36.35 | 37.2 | -0.8519 | 0.001125 |
| 21:26:35 | 36.34 | 37.2 | -0.8534 | 0.001087 |
| 21:31:35 | 36.37 | 37.23 | -0.8678 | 0.000753 |
| 21:36:35 | 36.4 | 37.24 | -0.836 | 0.001682 |
| 21:41:35 | 36.38 | 37.21 | -0.8318 | 0.001866 |
| 21:46:34 | 36.41 | 37.14 | -0.735 | 0.01859 |
| 21:51:34 | 36.41 | 37.1 | -0.6946 | 0.044589 |

Data compared by multiple t tests corrected for multiple comparisons using the Holm-Sidak method.

**Supplementary Table 2 - Primer sequences used for gene expression analysis and optimal conditions for qRT-PCR.**

| GENE | Primer Sequence (5' → 3') | Annealing Temperatures |
|---|---|---|
| *Bmal1* * | F:AAATCCACAGGATAAGAGGG<br>R:ATAGTCCAGTGGAAGGAATG | 58 °C |
| *Per2* * | F:CTTTCACTGTAAGAAGGACG<br>R:CTGAGTGAAAGAATCTAAGCC | 58 °C |
| *Clock* * | F:AAGTGACTCATTAACCCCT<br>R:CTATGTGTGCGTTGTATAGTTC | 60 °C |
| *Cry* * | F:AGAAGGGATGAAGGTCTTTG<br>R:CTCTTAGGACAGGTAAATAACG | 60 °C |
| *Hprt* | F: CTTCCTCCTCAGACCGCTTT<br>R: TCATCGCTAATCACGACGCT | 56 °C |

*Primer sequences previously reported by our group [52]

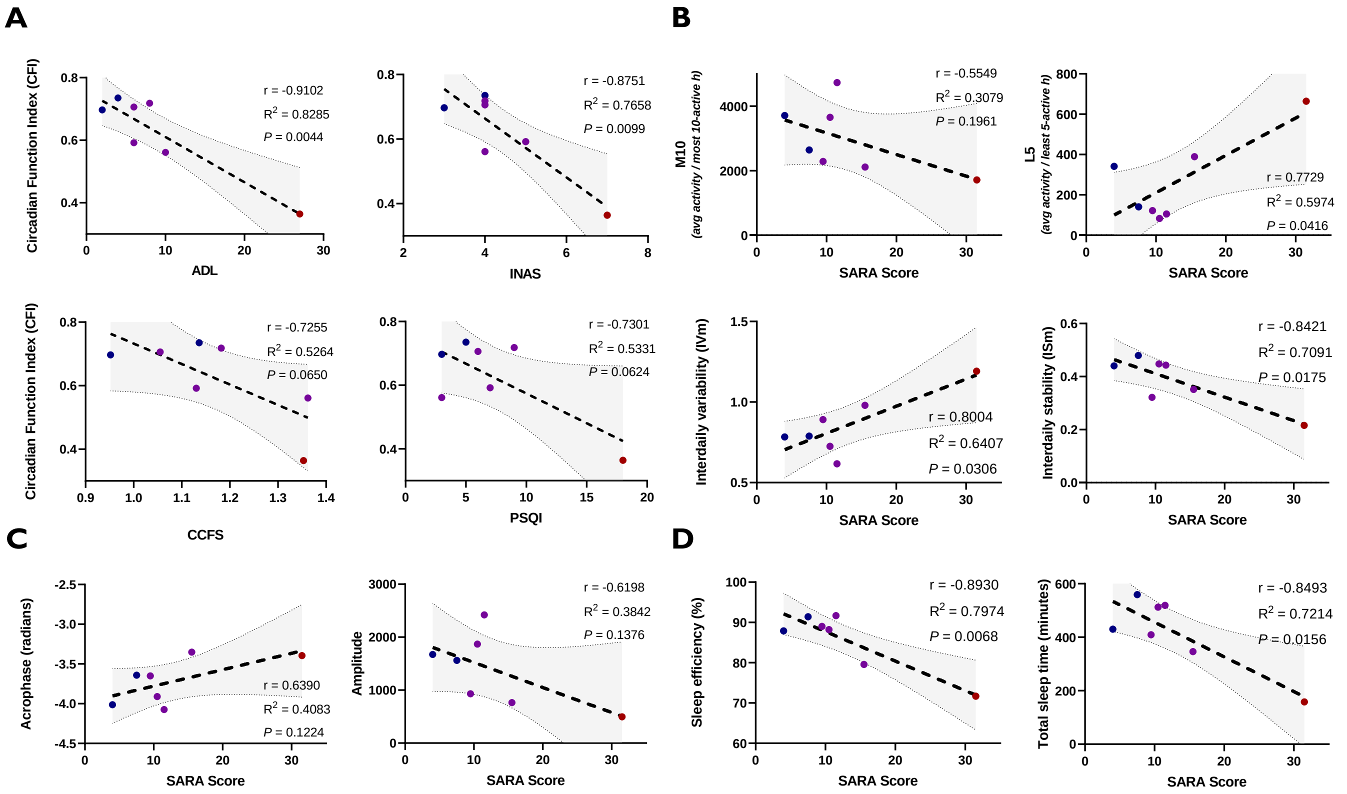


**Supplementary Figure 1 – MJD progression is associated with a decline of several circadian and sleep parameters (related to Figure 1).** (A) Negative correlations between Circadian Function Index (CFI) and the MJD clinical scales ADL, INAS, CCFS, and PSQI. (B) Circadian stability/synchronization (ISm) decreases, while rhythm fragmentation (IVm) increases with the development of the disease. M10 showed that activity during the day was not correlated with the progression; however, L5 suggests that activity increases during the rest phase despite the aggravated ataxia. (C) No significant differences found in the parametric measures acrophase and amplitude. (D) Sleep efficiency and total sleep time decreased with the progression of MJD given by the SARA score. Participants stratified by SARA score as mildly ataxic (3.0 ≤ SARA ≤ 7.5; dark blue), moderately ataxic (8.0 ≤ SARA ≤ 23.5; purple) or severely ataxic (SARA ≥ 24.0; red). Data distribution assessed using the Shapiro–Wilk test. All reported parameters showed data normally distributed and Pearson correlations were used to determine the correlation coefficient (r), goodness of the fit (R square), and *P* value. Graphs include 95% confidence bands of the best-fit line. Abbreviatures: Activities of Daily Living (ADL); Composite Cerebellar Functional Severity score (CCFS); Inventory of Non-Ataxic Signs (INAS); Pittsburgh Sleep Quality Index (PSQI); Scale for the Assessment and Rating of Ataxia (SARA).

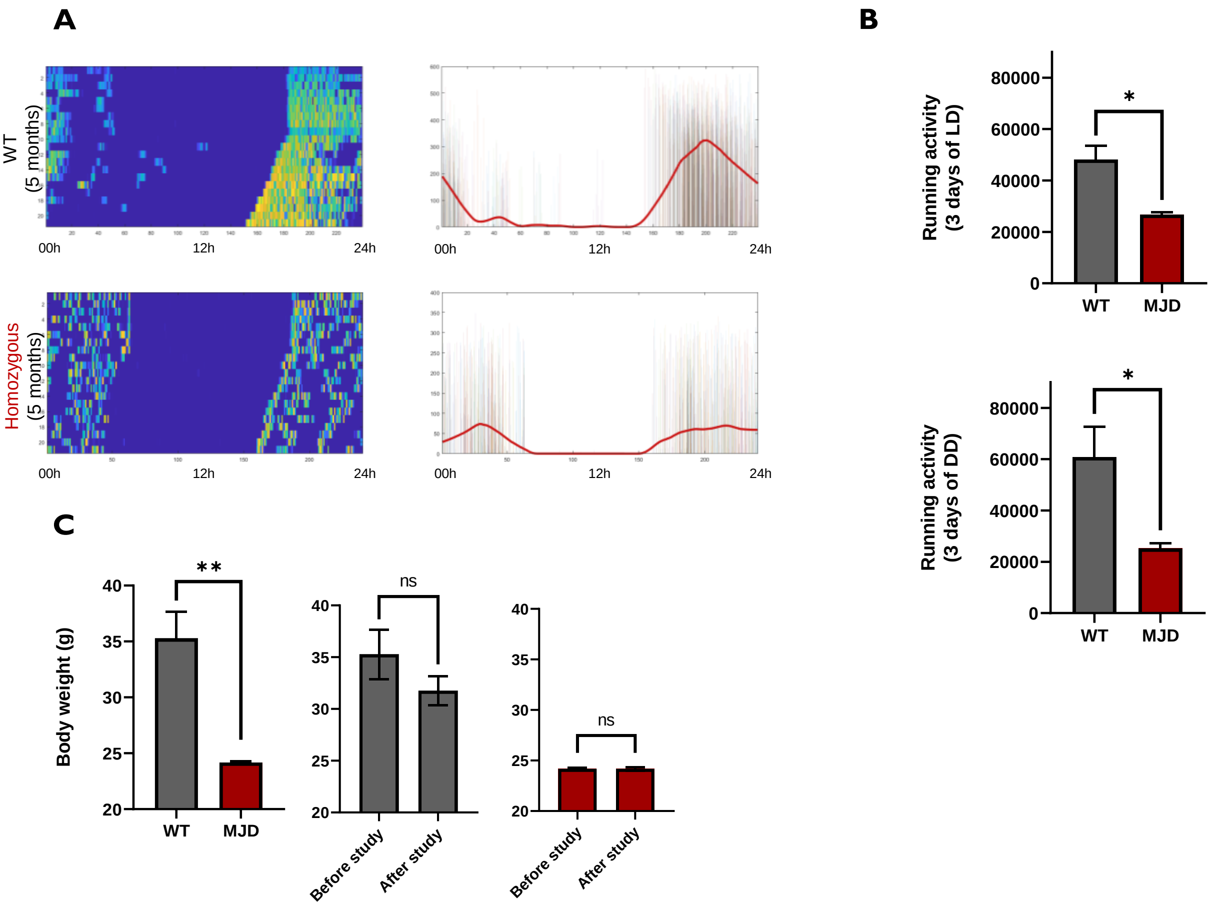


**Supplementary Figure 2 – YAC-MJD homozygous mice show less activity in both LD and DD periods and no significant changes in weight after the study (related to Figure 2).** (A-C) Data retrieved from the wheel-running study with MJD YAC84.2/84.2 (homozygous) and wild-type (WT) mice in constant darkness, after 8 days of LD entrainment. (A) MATLAB representation of the activity profile better elucidating fragmentation (left graphs) and activity differences per period of the day (right graphs). (B) Wheel revolutions in 3 complete days of LD (up) and DD (down), showing a decline in activity of the transgenic mice in both light schemes. (C) Weight differences between the homozygous mice and the WT control mice, characteristic of the YAC-MJD mouse model, and no significant variation of either the WT or homozygous weight after the activity study. Data is presented as mean ± SEM, include individual data (black dots), and groups are compared by unpaired t test (B, C) or paired t test for before and after the study comparisons (C). Statistical significance was set as: *$P < 0.05$, **$P < 0.01$. ns: not significant. Abbreviatures: light-dark (LD); dark-dark (DD).

**A**

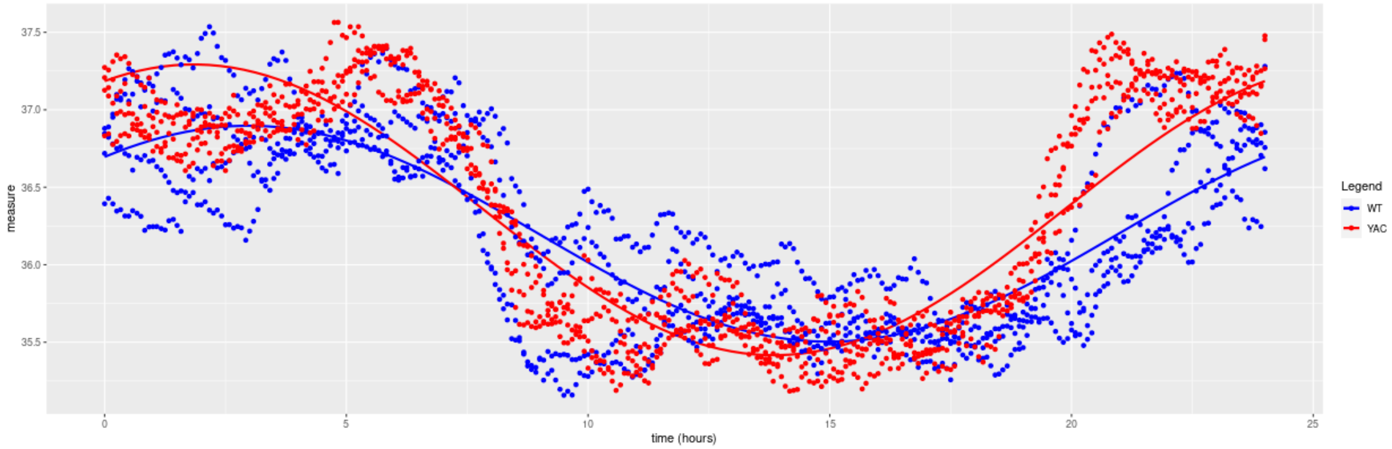


**Supplementary Figure 3 – CircaCompare graphical representation of average core body temperature values across the day (related to Figure 3).** (A) Graphical representation of mean core body temperature values calculated for every 5-minute timepoint across the 16 days of the MJD YAC84.2/84.2 and WT experiment (*n* = 4). Best fit curves were estimated based on MESOR, amplitude, and phase determinations using CircaCompare. Abbreviatures: wild-type (WT); Midline Estimating Statistic Of Rhythm (MESOR).

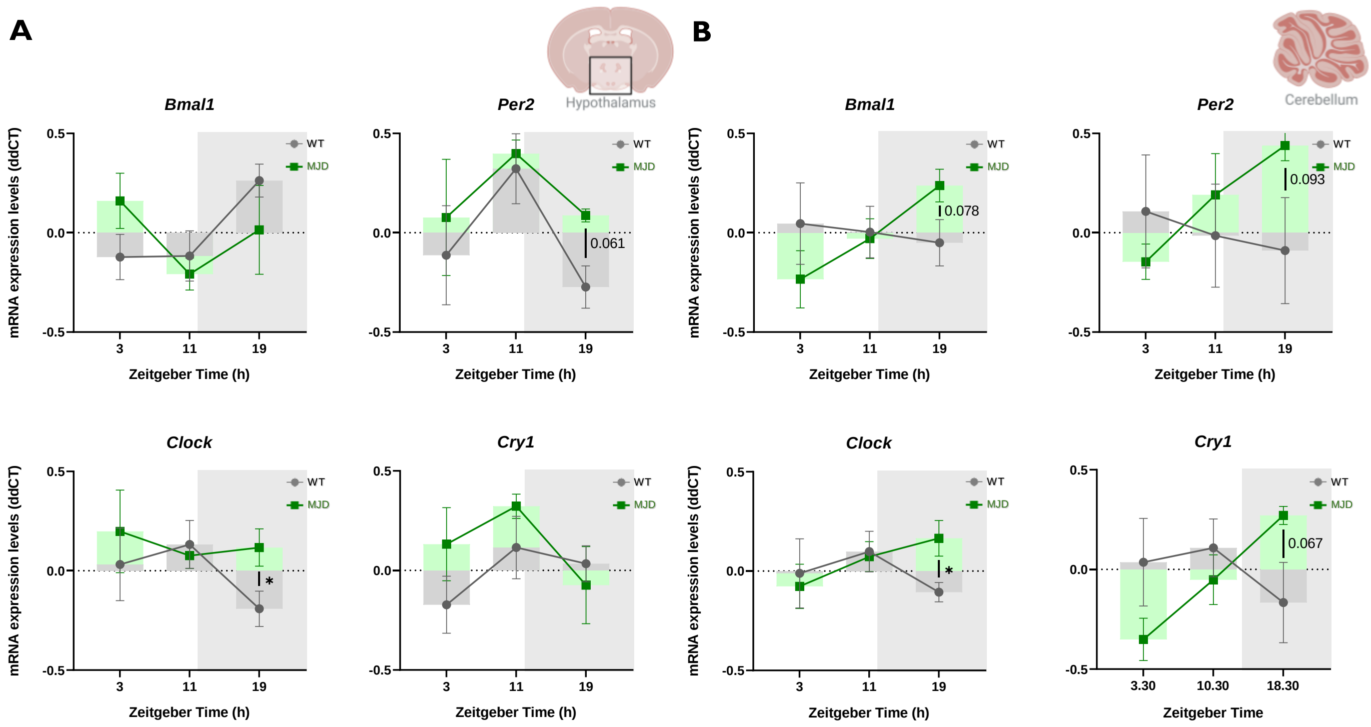


**Supplementary Figure 4 – YAC-MJD hemizygous mice show potential changes in clock gene expression, particularly in the cerebellum (related to Figure 5).** Expression levels of 4 core clock genes at 3 timepoints along the day analyzed in the (A) hypothalamus and (B) cerebellum of 10-12 month-old MJD YAC84.2 (hemizygous) and wild-type (WT) control mice. Despite the heterogenous population of the study (variable age, $n$ = 2-3 mice per timepoint/sex/genotype) the hemizygous mice showed tendencies for clock disruption, particularly in the cerebellum. Expression levels of target genes are shown relative to *Hprt*. Data presented as mean ΔΔCT values ± SEM. Grey rectangles indicate the dark phases. Unpaired t tests were performed to assess differences between hemizygous and WT control mice at each timepoint. Statistical significance set as: $*P < 0.05$.
